# How a highly acidic SH3 domain binds to its intrinsically disordered partner through the formation of an encounter complex intermediate

**DOI:** 10.64898/2026.07.28.741257

**Authors:** Valeria Jaramillo-Martinez, Ritika Kukreja, Michaela R. Cohen, Ally Mujica, Samuel Barton, Oluebube C. Onwuzulu, Jorge Cardoso, Matthew J. Dominguez, Isabelle M. Kekwick, Jaden Ali, Gemma M. Bell, Sydney Rice, Daniela Poaquiza, Colin McClure, Frida Anguiano, Michael P. Latham, K. Aurelia Ball, Elliott J. Stollar

## Abstract

Electrostatic interactions often play a role in determining the thermodynamic and kinetic properties of protein-protein interactions. However, the role of long-range electrostatic interactions in intrinsically disordered protein (IDP) binding is less clear, as they often bind in multiple steps including initial formation of a disordered encounter complex, followed by rearrangement into the bound state. We varied the salt concentration to probe the role of long-range electrostatic interactions in the binding of the highly charged AbpSH3 domain and the oppositely charged IDP ArkA. Using isothermal titration calorimetry, we observe that salt enthalpically destabilizes the bound complex. Molecular dynamics and NMR experiments reveal that salt has little effect on the bound state structure. However, simulations show that salt destabilizes the encounter complex intermediate, which primarily affects the association rate as measured by NMR. Consistent with these results, salt has the largest stabilizing effect on the apo SH3 domain, as cations substitute for the transient and long-range electrostatic interactions that can form with ArkA in the complex. We reveal a detailed picture of how a highly charged domain uses long-range, fuzzy, electrostatic interactions to help reach the bound state, a mechanism that is likely common among other highly charged domains that bind IDPs.

**TOC Image:** 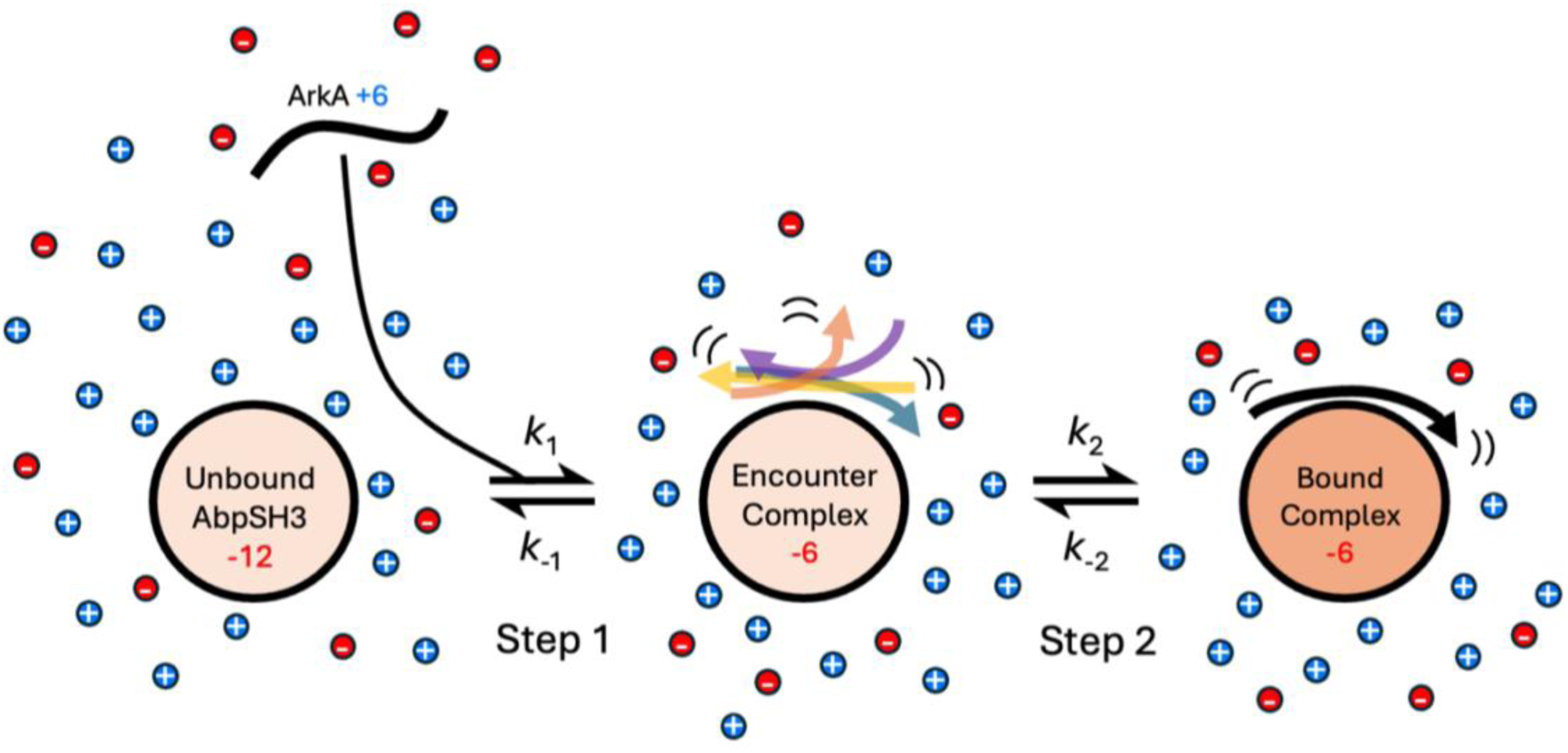

## 1. Introduction

Almost all proteins function through interactions with other proteins to form a variety of transient and obligatory complexes. Many protein-protein interactions involved in signal transduction involve partners that are disordered in the unbound state, allowing for multiple short-lived specific interactions, which are crucial for effective signaling. There is great interest in understanding the molecular mechanisms for these interactions involving intrinsically disordered regions (IDRs), which may not follow the same rules as binding interactions between folded domains due to their greater dynamics and flexibility^1,2^. One way that IDR binding partners may differ from folded partners is in the type of intermolecular interactions that control their binding kinetics and thermodynamics. Hydrogen bonding, van der Waals interactions, electrostatic interactions and hydrophobic packing all contribute to the stability of complexes, although their contributions to association and dissociation kinetics are hard to predict. Electrostatic interactions are the longest-range of these interactions, and therefore most easily probed by varying solution conditions. Increasing the ionic strength will disrupt long-range electrostatic interactions between binding partners, revealing the contribution of other shorter-range interactions to binding.

The clearest role of electrostatics in binding is when the binding partners have opposite net charge. At short range (less than 4 angstroms) attractive electrostatic interactions lead to specific salt-bridges and are the strongest non-covalent bond that exists in protein complexes. At long range (usually between 4 and 15 angstroms), attractive electrostatic interactions are weaker but can contribute to rapid protein-protein complex formation beyond the typical diffusion limit^3–5^. Furthermore, IDRs can form multiple long-range interactions with their target that can be present simultaneously in a less geometrically constrained manner to contribute to stabilization of the complex^6^. Charged residues are also found in proteins to increase solubility and prevent aggregation. In our previous study, we explored how an SH3 domain with high net-charge will fold despite the repulsion between like charges that must come together during the folding process^7^. Here we focus on how this highly-charged protein domain binds its oppositely-charged IDR binding partner.

The SH3 domain is one of the most common protein-protein interaction modules in eukaryotic cells, with around 300 encoded by the human genome alone^8^. Most SH3 domains are negatively charged and bind to disordered regions containing basic lysines and/or arginines^8^. We focus here on AbpSH3, the most negatively charged SH3 domain in *Saccharomyces cerevisiae*, with a net charge of negative twelve^7^. AbpSH3 has many binding partners, including Ark1p, involved in actin patch kinase localization and endocytosis^9^. AbpSH3 binds to the C-terminal IDR region of Ark1p, which contains 2 tandem PxxP binding sites called ArkA and ArkB. The ArkA binding site has been characterized previously and binds with higher affinity than ArkB and other known AbpSH3 partners, which also exhibit differential dynamic engagement with the SH3 domain^10^. Full-length ArkA binding site contains 17 residues (ArkA17) and six are positively charged lysines. Comparison with a shorter peptide, containing only three lysines (ArkA12), shows that binding affinity decreases six-fold when the net charge of the peptide is lowered^9^. However, hydrophobic interactions across the binding surface continue to stabilize the binding even when the net charge is decreased. The central lysine residue, K(-3), has been shown to be the most critical for binding, based on mutational studies and engages in both electrostatic and hydrophobic interactions with the AbpSH3 binding surface. Both the central and peripheral lysines also undergo conformational exchange in the bound state, while maintaining electrostatic interactions with negatively charged domain residues. Though electrostatics clearly play a role in the complex, the role of electrostatics in the binding pathway has not previously been probed directly.

Here, repeating the approach of our 2023 study on the AbpSH3 domain, we use salt to probe the effect of long-range electrostatic interactions, this time focusing on the binding process rather than folding. Salt principally interacts with charged proteins both through Debye screening and direct ion binding^11–13^. Both mechanisms will destabilize a complex formed between two oppositely charged proteins. This has been extensively studied in the case of protein-DNA binding^14^, as DNA is highly negatively charged, as well as for a complex that forms between two highly-charged disordered proteins^15^. Interactions involving intrinsically disordered regions (IDRs) are particularly sensitive to ionic strength due to their reliance on electrostatic forces. Changes in salt concentration can modulate these interactions by screening charges, thereby altering binding affinities and interaction specificity^15,16^. Meneses and Mittermaier have examined the effect of ionic strength on SH3 domain binding and found that while the association rate is decreased by salt due to electrostatic screening, this effect is not as pronounced as it is for more long-lived folded complexes such as Barnase and Barstar^3^. However, the effect of salt on the complete SH3 domain binding pathway that includes intermediates has not been examined in detail.

Our previous study on ArkA binding to AbpSH3 showed that by NMR, binding fits a two-state model with one transition state according to

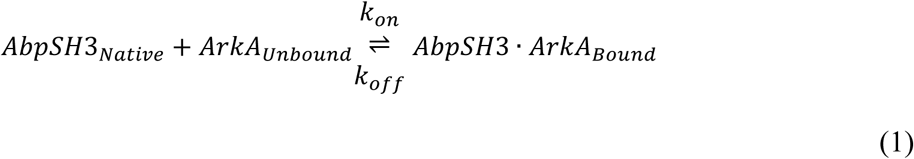

but molecular dynamics (MD) simulations reveal the formation of a transient encounter complex^17^. This loosely bound intermediate precedes the formation of the final, more ordered complex and plays a critical role in guiding productive binding. The existence of this intermediate supports the need for a two-step binding model, given by

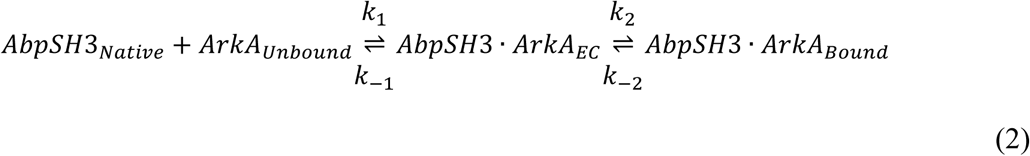

where an initial encounter complex forms rapidly, followed by a slower conformational rearrangement into the fully bound state. This fits the three-state “dock and coalesce” mechanism^5^, which better captures the underlying kinetics of IDP–domain interactions. In general, for protein-protein interactions, either step in a two-step process could be rate-limiting, but for IDP binding to a folded domain we expect the first step to be faster than the second, indicating that the rate-limiting step for binding occurs after the encounter complex formation. This is because the IDP is disordered in the encounter complex, and therefore not geometrically constrained, leading to a rapid initial association that is controlled by the diffusion rate (without a large entropic barrier), which can be enhanced by long-range electrostatics. Thus, the second “coalesce” step of the process, involving slower stereo-specific rearrangements, will be slower than the “docking” step. To address the complexities involved in two-step IDP binding enhanced by electrostatic interactions, we use a combined approach to examine the effects of salt, employing isothermal titration calorimetry (ITC) to probe binding thermodynamics, NMR to probe dynamics and binding kinetics, and MD simulations to probe the encounter complex intermediate and atomistic interactions. Together, we are able to form a complete picture of how salt affects the binding pathway and therefore how electrostatic interactions contribute to binding.

## 2. Methods

### 2.1 Plasmids and proteins

For ITC and NMR, unlabelled AbpSH3 domain was required and was expressed in BL21 (DE3) Gold *E.coli* cells as an N-terminal Deca-Histidine tagged protein within the pET21 vector with a T7 promoter and ampicillin resistance (synthesized by Twist Biosciences) with the sequence MHHHHHHHHHHPKENPWATAEYDYDAAEDNELTFVENDKIINIEFVDDDWWLGELEKDGSKGLFPSNYVSLGN. Protein was purified by a denaturing IMAC method on a 20 mL column with 5 mL Ni-NTA beads as previously described^18^ and subsequently purified using anion exchange chromatography with Q Sepharose Fast Flow beads (GE healthcare, UK). Eluates from 0.2-0.6 M NaCl were pooled and dialysed into 10 mM Tris, pH 8.1 and concentrated using Vivaspin 20, 3kDa MWCO columns (GE healthcare, UK) following the manufacturer’s protocol.

For NMR backbone dynamics measurements, ^15^N-labelled ArkA17 peptide was required and expressed in BL21 (DE3) Gold *E.coli* cells as a Thioredoxin fusion in 1 L in minimal media (supplemented with ^15^N-labelled ammonium chloride) and purified by denaturing IMAC method on a 20 mL column with 5 mL Ni-NTA beads as previously described (Stollar 2009). The sample was cleaved with TEV protease with an enzyme to protein molar ratio of 1:32 in 20 mM Tris, pH 8, 300 mM NaCl, 5 mM DTT, 0.5 mM EDTA, and was left overnight at 4 °C and for 3 hours the next day at room temperature. To separate the domain from the peptide, the sample was diluted 5-fold in 20 mM Tris, pH 8 and further purified on 0.25 mL SP Sepharose Fast Flow cation exchange resin (GE Healthcare) column and eluted in increasing concentrations of Ammonium bicarbonate (NH_4_)HCO_3_ pH 8, from 50 to 1500 mm with elution fractions (150 mM -250 mM) containing peptide pooled and lyophilised. The final NMR sample contained 500 µM ^15^N-ArkA17 with 1.2 mM unlabeled AbpSH3 in 10 mM Tris, pH 8.1, 1X protease inhibitor cocktail (Cambridge Biosciences), 1mM EDTA, 1 mM Sodium Azide and either 0, 0.1 M, 0.5 M or 0.8 M NaCl.

For NMR CPMG and HSQC experiments, the ^15^N-labelled AbpSH3 domain was expressed NiCo21 (DE3) competent *E.coli* cells in minimal media and expression was induced with IPTG. The ^15^N-labelled AbpSH3 domain was purified using a denaturing purification using nickel affinity chromatography^18^. This was followed by anion exchange chromatography, and the pure AbpSH3 domain was dialyzed into 10 mM Tris buffer for NMR. Further details of expression and purification are provided in the supporting information.

Unlabeled synthetic peptides required for NMR and ITC were synthesized with an N-terminal acetyl group and a C-terminal amide group to neutralize the termini, purified and desalted by Peptide 2.0. The CPMG NMR samples contained 1 mM ^15^N-labeled AbpSH3 and 70 μM peptide (ArkA17, ArkA12 or ArkA10GS) in 10 mM Tris, pH 8.1, 0.1 mg/mL AEBSF protease inhibitor (Thermo Scientific), 1 mM EDTA, 1 mM Sodium Azide and either 0, 0.1 M, 0.5 M or 0.8 M NaCl. The HSQC samples contained 0.1 mM ^15^N-labeled AbpSH3 and excess (1 mM) peptide (ArkA17, ArkA12 or ArkA10GS) in 10 mM Tris, pH 8.1, 0.1 mg/mL AEBSF protease inhibitor (Thermo Scientific), 1 mM EDTA, 1 mM Sodium Azide and either 0, 0.1 M, 0.5 M or 0.8 M NaCl.

Domain-peptide hybrids were expressed in BL21 (DE3) *E.coli* cells as an N-terminal Deca-Histidine tagged protein within the pET21 vector with a T7 promoter and ampicillin resistance (synthesized by Twist Biosciences). The sequence of the AbpSH3-ArkA17 hybrid was as follows (both the His-tag region and the linker between the domain and peptide as shown in bold): **MQLSHHHHHHHHHHLEVLFQGPGTS**NPWATAEYDYDAAEDNELTFVENDKIINIEFV DDDWWLGELEKDGSKGLFPSNYVSLGN**GSENLYFQGGSGT**KKTKPTPPPKPSHLKPK. A starter culture in TB supplemented with 100 ug/mL of carbenicillin was grown overnight and used to inoculate in 100 mL autoinduction media (Formedium) and grown at 37°C during the day and then 20°C overnight. Protein was purified by a native IMAC method on a 20 mL column with 5 mL Ni-NTA beads as previously described^18^, except cells were lysed using (4 mL per 1 g wet weight cells) of B-PER reagent supplemented 1 mM EDTA as per the manufacturer’s instructions.

### 2.2 Isothermal titration calorimetry

Isothermal titration calorimetry (ITC) experiments were performed using a MicroCal PEAQ-ITC instrument (Malvern). Binding of the SH3 domain to synthetic peptides ArkA17, ArkA12, ArkA10 were measured in 10 mM Tris, pH 8.1, supplemented with 0, 100, 500, or 800 mM NaCl, at a constant temperature of 30 °C.

Protein and peptide samples were prepared in matching buffers. Peptides were weighed, dissolved to 50 mg mL⁻¹, clarified by centrifugation, and quantified spectrophotometrically at 214 nm using extinction coefficients of 36350, 28896 and 33624 M^-1^ cm^-1^ respectively based on calculations by Kuipers and Gruppen^19^. SH3 domain samples were clarified (max speed, 5 min, 4 °C), quantified, and diluted to the desired concentrations. All buffers and samples were degassed immediately prior to loading.

ITC titrations were carried out with the SH3 domain placed in the cell and peptide in the syringe. Experimental conditions were as indicated in Table 1.

**Table 1.** ITC titration details.

| NaCl concentration (mM) | SH3 domain concentration in the cell (μM) | Peptide concentration in the syringe (μM) | Number of injections |
| --- | --- | --- | --- |
| 0 | 10 | 100 | 19 |
| 100 | 20 | 250 | 13 |
| 500 | 40 | 400 | 13 |
| 800 | 40 | 400 | 13 |

Stirring speed and reference power were kept at instrument defaults. Raw thermograms were integrated and fitted using the MicroCal PEAQ-ITC Analysis software. A one-site binding model was applied, and the stoichiometry was set to 1 in the fitting process, allowing the peptide concentration to change within 20% of its measured value. Heats of dilution were subtracted using control injections of peptide into buffer.

### 2.3 NanoDSF on Prometheus Panta

10 μM samples of the AbpSH3-ArkA hybrids and AbpSH3 alone were prepared in 10 mM Tris, pH 8.1, supplemented with 0, 100, 500, or 800 mM NaCl and underwent reversible thermal denaturation using a 1 °C per minute temperature ramp monitoring tryptophan fluorescence emission at 330 and 350 nm. The major inflection point within the melting curve was determined by taking the derivative of the melting curves of temperature versus the 350/330 nm ratio. Experiments were performed in triplicate.

### 2.4 NMR spectroscopy

NMR data was collected at 25 °C (for ^15^N labeled AbpSH3) and 10 °C (for ^15^N labeled ArkA peptide) using Bruker Avance III 900 MHz (21.14 T), Bruker 850 MHz Avance III (19.97 T), and Bruker 600 MHz Avance NEO (14.1 T) spectrometers equipped with 5 mm TCI cryoprobe with z-axis gradient. NMR data were processed with NMRPipe/NMRDraw^20^ and analyzed with CCPN analysis v2.5^21^. 2D ^15^N,^1^H HSQC spectra were collected for both ^15^N AbpSH3 domain bound to 5-10% unlabeled peptide and ^15^N ArkA peptide bound to saturated unlabeled AbpSH3 domain (1:2) at 0 M, 100 mM, 500 mM and 800 mM NaCl. Backbone amide ^1^H and ^15^N assignments for Apo (unbound) AbpSH3 domain and Apo ArkA peptide were taken from previous studies^7^. Amide chemical shift perturbations (CSPs) were calculated as

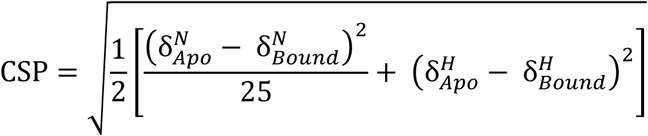

where *δ^N^*and *δ^H^* are the nitrogen and proton chemical shifts, respectively.

Backbone ^15^N R_1_ and R_1ρ_ and ¹⁵N-{^1^H} heteronuclear NOE experiments were performed for ^15^N ArkA bound to unlabeled AbpSH3 domain at 600 MHz at 10 °C (500 µM ^15^N ArkA with 1.2 mM unlabeled AbpSH3). ^15^N R_1_ and R_1ρ_ relaxation rates were calculated from the single exponential fits of the intensities from six and seven parametrically varied time points ranging from 2– 200 msec (R_1_) and 2–60 msec (R_1ρ_), respectively. R_1ρ v_alues were converted to R_2_ using the relation

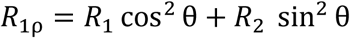

where *θ*=*tan*^−1^(*ν_SL_*/Ω) is the effective tilt angle of the rotating frame, *ν_SL_* is the field strength of the applied spin lock (2000 Hz), and Ω is the offset of the peak from the ^15^N carrier. Errors in the relaxation rates were calculated from the covariance matrix of the fit. Backbone ^15^N R_1_, R_2_, and NOE data were used to calculate residue-specific order parameters (S^2^) and the overall global rotational correlation time (τ_c_) using the model-free approach as implemented in Modelfree v4.2 (https://comdnmr.nysbc.org/comd-nmr-dissem/comd-nmr-software/software/modelfree). All data were fit using “model 2” using an axially symmetric diffusion model resulting in an average τ_c_ value of approximately 7.92 ± 0.24 ns across the salt concentrations. NH bond lengths of 1.02 Å and ^15^N chemical shift anisotropy of −160 ppm were used in the fitting.

¹⁵N CPMG relaxation dispersion profiles were recorded for ^15^N AbpSH3 bound to 5-10% unlabeled peptide at four different salt conditions (0 M,100 mM, 500 mM and 800 mM NaCl) at ¹H Larmor frequencies of 600 MHz (14.09 T) and 900 MHz (21.14 T) using the sensitivity-enhanced, CW-decoupled pulse sequence of Hansen et al., with a 40 ms constant-time relaxation delay at both fields^22^. A series of 16 CPMG frequencies ranging from 12.5 to 1000.0 Hz were sampled by varying the number of refocusing cycles within the constant-time delay, with two frequencies repeated for error estimation. A reference spectrum with no CPMG refocusing was also acquired at each field. Dispersion data were fitted using the program ChemEx (https://gbouvignies.github.io/ChemEx/) to a two-site global exchange model by numerically solving the Bloch-McConnell equations^22–24^.

### 2.5 MD simulations

#### 2.5.1 Simulations

To determine the effect of salt on the ArkA-AbpSH3 interactions at atomic resolution, MD simulations of the bound complex and the binding process were run at high salt concentrations. For the bound complex, we ran twenty independent simulations each for the ArkA12 and ArkA17 bound complexes, ten at 0 mM sodium chloride and ten at ∼900 mM sodium chloride (Table 2). These simulations were initiated from the lowest energy NMR structure of ArkA17 bound to AbpSH3 (PDB: 2RPN)^9^. For the ArkA12 simulations, the ArkA17 residues at the N and C termini were removed and replaced with a capping acetyl group (ACE) or amide group (NHE) respectively. Most bound simulations were run for 2.4 μs except for the ArkA17 bound simulations in high salt, which were run for 900 ns. To simulate the binding process and the encounter complex, ArkA12 binding simulations were run as in Gerlach et al.^17^, but with ∼800 mM sodium chloride in the simulation box (Table 2). Due to the small box size and the differing number of sodium and chloride ions needed to keep the system neutral (Table 2), the exact NaCl concentrations are different whether sodium or chloride ions are used to calculate concentration. Therefore, we refer to the NaCl concentration in the simulations based on the approximate concentration rounded to the nearest 100 mM. Previously reported simulations of ArkA12 binding in 0 mM salt (counter ions only) and of the AbpSH3 domain alone in high salt were also compared^7,17^. Table 2 indicates the details of each of the different simulations compared in this paper.

**Table 2.**
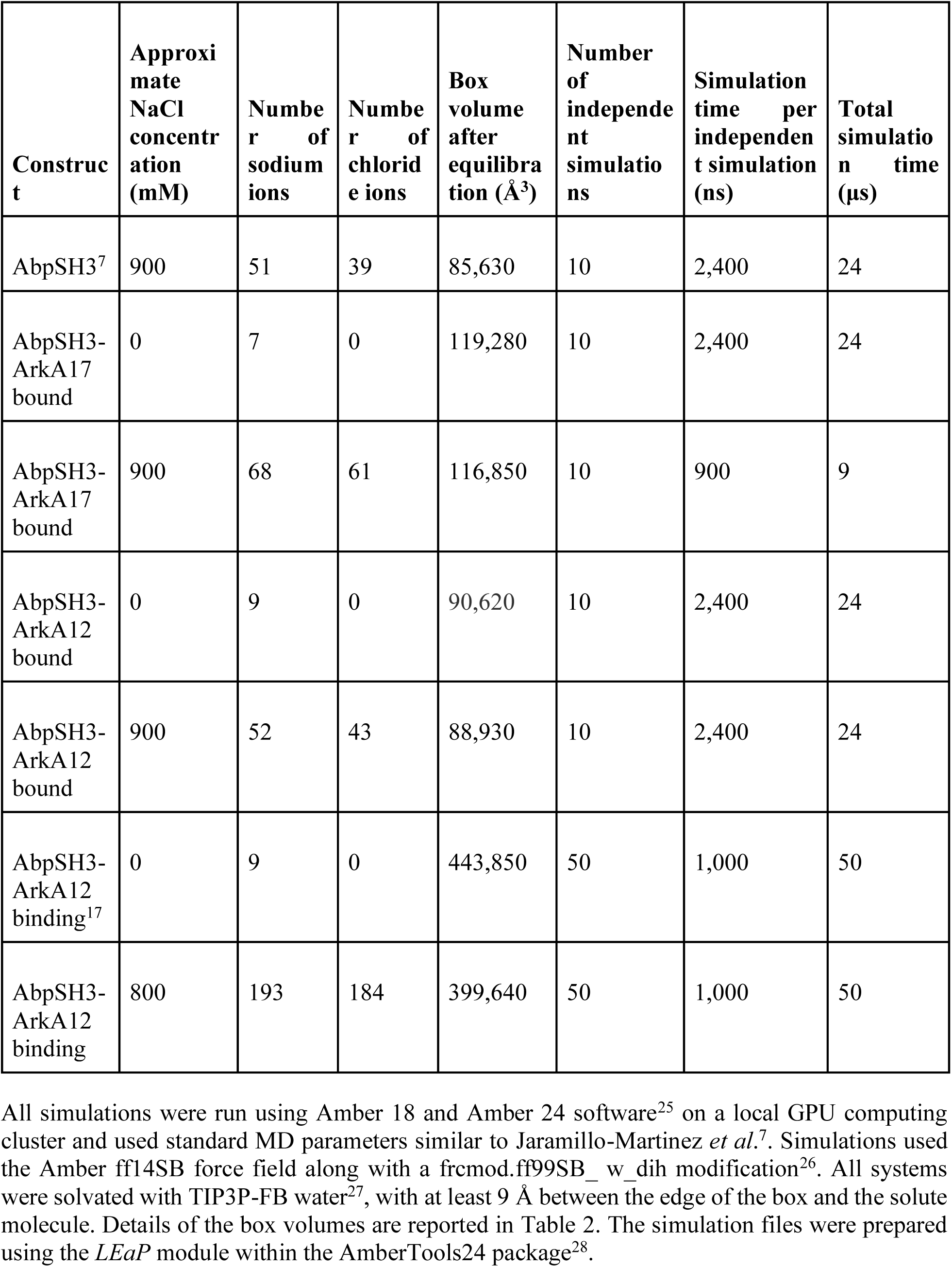
MD simulation details.

| **Construct** | **Approximate NaCl concentration (mM)** | **Number of sodium ions** | **Number of chloride ions** | **Box volume after equilibration ($\text{\AA}^3$)** | **Number of independent simulations** | **Simulation time per independent simulation (ns)** | **Total simulation time ($\mu\text{s}$)** |
| --- | --- | --- | --- | --- | --- | --- | --- |
| AbpSH3<sup>7</sup> | 900 | 51 | 39 | 85,630 | 10 | 2,400 | 24 |
| AbpSH3-ArkA17 bound | 0 | 7 | 0 | 119,280 | 10 | 2,400 | 24 |
| AbpSH3-ArkA17 bound | 900 | 68 | 61 | 116,850 | 10 | 900 | 9 |
| AbpSH3-ArkA12 bound | 0 | 9 | 0 | 90,620 | 10 | 2,400 | 24 |
| AbpSH3-ArkA12 bound | 900 | 52 | 43 | 88,930 | 10 | 2,400 | 24 |
| AbpSH3-ArkA12 binding<sup>17</sup> | 0 | 9 | 0 | 443,850 | 50 | 1,000 | 50 |
| AbpSH3-ArkA12 binding | 800 | 193 | 184 | 399,640 | 50 | 1,000 | 50 |

All simulations were run using Amber 18 and Amber 24 software^25^ on a local GPU computing cluster and used standard MD parameters similar to Jaramillo-Martinez *et al*.^7^. Simulations used the Amber ff14SB force field along with a frcmod.ff99SB_ w_dih modification^26^. All systems were solvated with TIP3P-FB water^27^, with at least 9 Å between the edge of the box and the solute molecule. Details of the box volumes are reported in Table 2. The simulation files were prepared using the *LEaP* module within the AmberTools24 package^28^.

Two rounds of minimization were performed, the first restraining the protein with harmonic restraints (10 kcal/mol) and the second without restraints. Each round consisted of 500 cycles of steepest descent minimization and 500 cycles of conjugate gradient minimization. After minimization, a simulation with restraints on the protein was performed to heat the system to the final desired temperature of 300K over 40 ps. Next, the system was equilibrated at constant temperature and pressure using two sequential equilibration steps, the first with restraints on the protein, and the second unrestrained. The restrained equilibration step lasted 50 ps and used a Berendsen barostat. The unrestrained equilibration phase lasted 200 ps and used the Monte Carlo barostat. Temperature was controlled using a Langevin thermostat. During both phases of equilibration, periodic boundary conditions were applied, and the Particle Mesh Ewald procedure was used to handle long-range electrostatics with a non-bonded cutoff of 9 Å for the direct space sum. Independent production simulations were then run using random starting velocities. Production simulations were run in the NPT ensemble at 300 K and 1.013 bar, using Langevin dynamics with a collision frequency of 1.0 ps^-1^, with a Monte Carlo barostat, new system volumes attempted every 100 steps, an integration step every 2 fs, and coordinates stored every 10 ps. Bonds to hydrogen were constrained using the SHAKE algorithm. Table 2 indicates the number of independent simulation runs for each system and the total simulation time for each system.

#### 2.5.2 Analysis

The *cpptraj* module of AmberTools24 was used in combination with in-house python scripts for all MD simulation analysis including atomic fluctuations, distance measurements, solvent accessible surface area, and hydrogen bond analysis^28^. Most analysis was performed on structures stored every 10 ps from production simulations. Error bars were calculated by taking the standard error across independent simulations. To compare simulations with and without salt present, *p*-values for the means were calculated using a permutation test. All *p*-values were found using 10,000 permutations. Structural figures were created with Visual MD^29^.

The binding surface distance was calculated as in Gerlach et al. 2020 to quantify the extent of binding^17^, and is described in the supporting information (Figure S1). Snapshots with a binding surface distance below 11.5 Å were designated as bound, distances between 11.5 Å and 23 Å were labeled as encounter complex, and above 23 Å the system was considered unbound. Dihedral angle RMSD, solvent accessible surface area (SASA), and specific hydrophobic interactions were calculated as in Gerlach et al. 2020^17^.

Contact maps were created using the open-source Contact Map Explorer python package with a cutoff of 4.5 nm for any two heavy atoms. For electrostatic and ion contacts, the distance was measured from the positively charged nitrogen atom on the lysine side chain and the most distal carbon atom of the glutamate or aspartate side chain. These distance measurements were used to calculate the percent of time that electrostatic and ion contacts were present in the simulations. A cutoff distance of 3 Å was used for ion contacts (Figure S7), 4 Å between the terminal carbon or nitrogen atom on the charged side chain was used for short-range electrostatic contacts and residue contacts with ions, and 10 Å was used for long-range electrostatic contacts. Only interactions present in at least one of the simulations for at least 10% of the ensemble were included in the results.

The contact map differences between simulations with and without salt were compared using two methods; one comparing the average number of contacts, and one comparing the sum of squared contact population differences. The average number of contacts for each dataset was found by summing the percentage values within the contact maps in each independent simulation. Using the method of the sum of squared contact population differences, first a test statistic (the value presented in Table S2) was calculated from the cells of each contact population difference cell summed together. The p-value for this test statistic was determined using a permutation test, finding the proportion of the 10,000 statistics that were greater than the original test statistic. Though comparing the average number of contacts provided an important metric to determine how different the datasets with and without salt are, our comparison of the sum of squared contact population differences provides an additional metric that can be used in our analysis of how much salt affects contacts made between residues in ArkA and AbpSH3.

Binding rate constants were calculated using a Bayesian inference method with a Jeffreys Prior^30^. The point estimate of the binding rate (*k*_1_ or *k_on_*) is given by

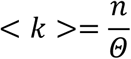

where *n* is the number of independent simulations that bind and *Θ* is the total simulation time before binding which includes the full simulation time for simulations that never bound and the time before the first binding event for those that did bind. The variance on this point estimate of the rate constant is given by

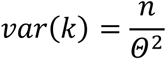

To determine if salt had an effect on the rate constants in our simulations, we calculated the odds ratio using equation 38 from Ensign and Pande^30^,

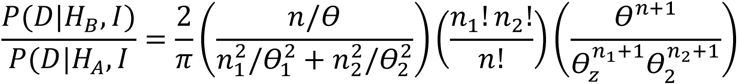

A pseudo-dissociation constant 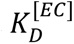 for the encounter complex was calculated as the ratio of the dissociation rate for the encounter complex over the encounter complex association rate,

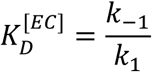

We also calculated 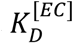 based on populations in our MD simulations of binding for comparison, by taking the ratio of the total time spent in the unbound state over total time spent in the encounter complex,

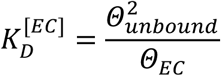

We considered the unbound state and encounter complex to be in pseudo-equilibrium in our simulations as we observed many transitions between those two states during our simulation time. To remove the bias from starting all of the simulations in the unbound state, we calculated the time *t*_1/2_ at which half of our binding simulations had reached the encounter complex and only included the time spent after *t*_1/2_ in our calculation of *Θ_unbound_* and *Θ_EC_* . This value for 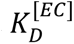 is reported in the supporting information (Table S4).

## 3. Results and Discussion

### 3.1 Long-range electrostatic attraction stabilizes the AbpSH3-ArkA complex

Multiple attractive long-range electrostatic interactions can form when the negatively charged AbpSH3 is bound to its positively charged target peptide, ArkA17 (Figure 1A). Isothermal Titration Calorimetry (ITC) was used to probe the electrostatic contribution to binding for ArkA17 as well as two variant peptides with different net positive charge (Table 3). The binding free energy *ΔG_N_*_→*B*_ (going from the native unbound state to the peptide-bound state) decreases with the square root of ionic strength for all three peptides, showing that salt decreases the binding affinity, as predicted by the Debye-Hückel rate limiting law (Figure 1B). ITC data was consistent with melting temperature data with these peptides within a AbpSH3-peptide hybrid, where the peptide is attached the C-terminus of the domain via a flexible linker (Figure S2). This is analogous to the increase in AbpSH3 stability with ionic strength that we have previously reported^7^. As expected, the binding strength in low salt conditions follows the net charge of the peptide, with ArkA17 showing the largest change in binding affinity with salt and ArkA10GS the smallest. At 800 mM salt, there is little difference in binding affinities for the three peptides, suggesting that most of the long-range electrostatic interactions have been screened and the peripheral lysine residues have little contribution to binding affinity under these conditions. The central lysine at position -3, which is known to form a critical salt bridge^9^, is likely still playing a role even at high salt concentrations, as this short-range interaction is comparable to the Debye-Hückel screening length of ∼3.4 Angstroms at 800 mM NaCl^31^.

**Fig 1:**
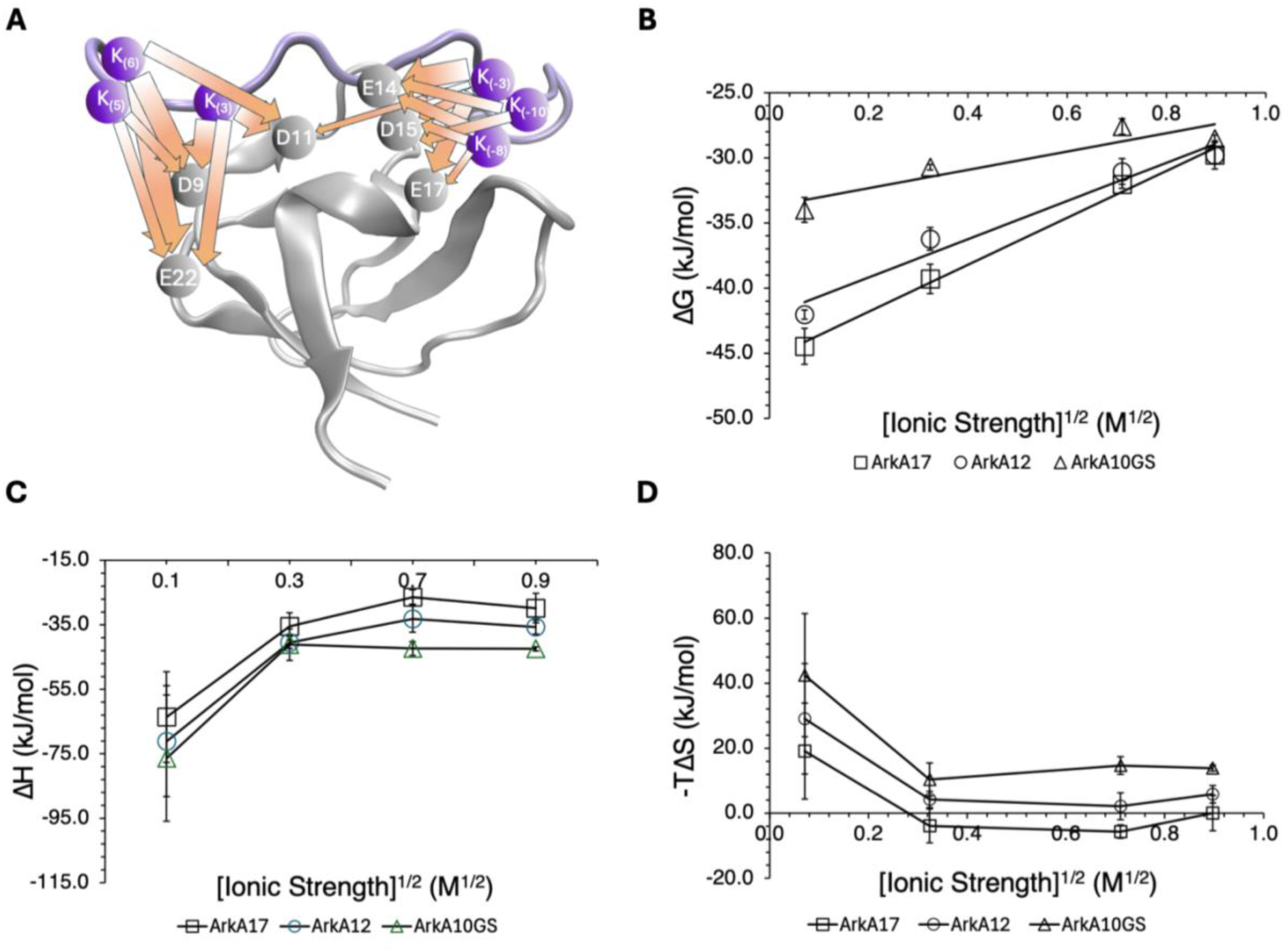
**AbpSH3-ArkA binding affinity decreases with ionic strength**. (A) Visual representation of the intermolecular long-range electrostatic interactions from simulations. Arrows represent the interactions that are present in at least 10% of the ensemble from the AbpSH3-ArkA17 simulations without salt, and the arrow thickness is based on the frequency of the interaction. Long-range electrostatic interactions are considered formed when the two charged groups are within 10 Å of each other. (B) Binding free energy measured by ITC plotted versus the square root of ionic strength for ArkA17, ArkA12, ArkA10GS with R^2^ values of 0.85, 0.96 and 0.99 respectively. Error bars represent standard deviations across experiment repetitions (some of the error bars are smaller than the markers with values less than ±2%). (C) Enthalpy of binding measured by ITC with varying ionic strength for ArkA17, ArkA12, and ArkA10GS. (D) Entropy of binding calculated based on the binding free energy and enthalpy of binding from ITC with varying ionic strength for ArkA17, ArkA12, and ArkA10GS.

**Table 3.** ArkA peptides.

| Peptide | Net charge | Sequence |  |  |  |  |  |  |  |  |  |  |  |  |  |  |  |  |
| --- | --- | --- | --- | --- | --- | --- | --- | --- | --- | --- | --- | --- | --- | --- | --- | --- | --- | --- |
|  |  | 6 | 5 | 4 | 3 | 2 | 1 | 0 | -1 | -2 | -3 | -4 | -5 | -6 | -7 | -8 | -9 | -10 |
| ArkA17 | +6 | K | K | T | K | P | T | P | P | P | K | P | S | H | L | K | P | K |
| ArkA12 | +3 |  |  |  | K | P | T | P | P | P | K | P | S | H | L | K |  |  |
| ArkA10G<br>S | +1 | G | S | G | A | P | T | P | P | P | K | P | S | H | L | A | G | S |

Salt could be affecting binding in two ways. It could break up interactions in the bound state by replacing interactions between the two proteins; this would make it easier for the peptide to dissociate from the domain and therefore decrease the barrier to dissociation, *ΔG_B_*_→†_ (the free energy to go from the bound state to the transition state). Additionally, it could form favorable interactions with the unbound domain and free peptide, this would make it harder for the peptide to associate with the domain and therefore increase the barrier to association, *ΔG_N_*_→†_ (the free energy to go from the unbound state to the transition state). The previous result that salt stabilizes the folded domain in the absence of peptide predicts a salt effect on *ΔG_N_*_→†_ (Jaramillo-Martinez 2023). However, there may also be an effect on *ΔG_B_*_→†_ . To determine how salt affects each of these barriers, we use complimentary techniques from NMR and MD simulations.

### 3.2 Bound peptide dynamics and structure are minimally affected by salt

To evaluate potential changes in the structure of the ArkA17–AbpSH3 complex as a function of ionic strength, we acquired 2D ¹⁵N-{^1^H} correlation NMR spectra of ¹⁵N-labeled AbpSH3 with saturating concentrations of unlabeled ArkA17, ArkA12, or ArkA10GS (Figure S3A). The increased salt concentration has modest effects on the chemical shift changes between free domain and peptide-bound domain (Figure S3B), confirming the structure of the complex is similar at the different salts. We also measured 2D ¹H-¹⁵N correlation NMR spectra of ¹⁵N-labeled ArkA17 in the presence and absence of saturating concentrations of unlabeled AbpSH3. The spectrum of the labeled peptide bound to the unlabeled domain exhibited well-dispersed resonances consistent with the peptide in the fully bound state, and only minor chemical shift perturbations were observed across salt concentrations, indicating that elevated ionic strength has negligible effects on the overall structure of the bound complex (Figure 2A). Several residues peripheral to the binding interface: T(4), T(1), and S(−5), exhibited progressive line-broadening with increasing salt concentration, which we also observed for several residues in the labeled domain spectrum (Figure S3A). Given that this broadening occurred for residues across the peptide and domain beyond the binding surface, we attribute this to a general effect of salt on the exchange regime of the local chemical environment of the protein backbone and not to any change in the way that the peptide and domain interact in the bound state. Overall, the HSQC spectra show that the addition of salt causes minimal changes to the structure of complex.

**Fig 2:**
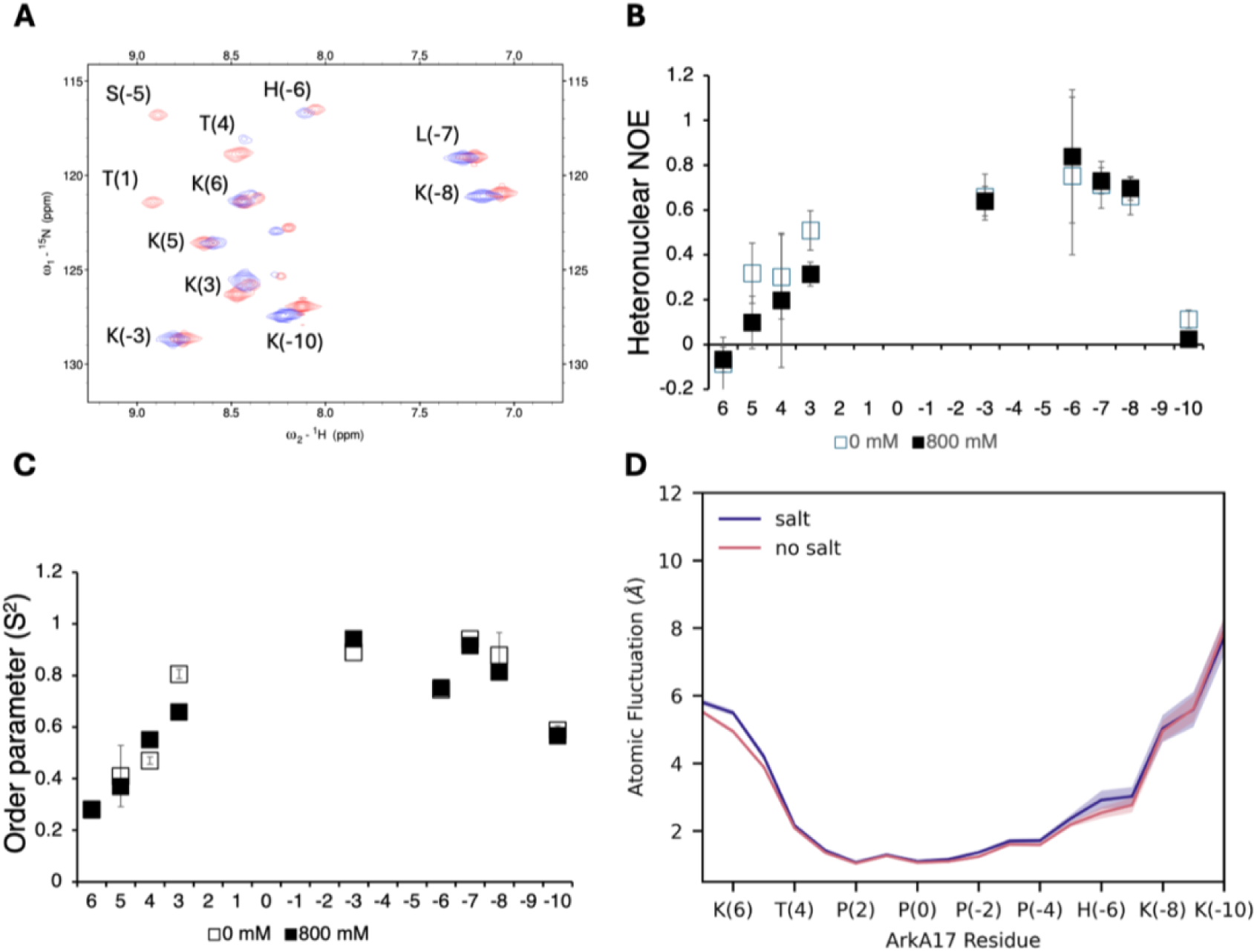
**Bound ArkA shows little change in fast-timescale dynamics and structure with increasing salt concentrations**. (A) Bound ¹⁵N-labeled ArkA17 HSQC overlay at two salt concentrations, 0 mM (red) and 800 mM (blue). Peaks are assigned to ArkA residues, but no proline residues are present because of the lack of amide proton in the proline backbone. The two small peaks that are not labeled are from N-terminal residues that are there due to the cloning process but not part of the ArkA motif. (B) Heteronuclear NOE of bound ArkA17 at two salt concentrations, 0 mM (open squares) and 800 mM (filled squares) (C) Order parameter of bound ArkA17 at two salt concentrations, 0 mM (open squares) and 800 mM (filled squares). For residues K(6) and H(-6), the 0 mM marker is completely obscured by the 800 mM marker. Residues T(1) and S(-5) are not present in the NOE and order parameter plots because those peaks are missing in the 800 mM NaCl spectrum. (D) Root-mean-squared back bone fluctuations of ArkA17 bound to AbpSH3 from simulations at 0 mM (pink) and 900 mM (blue) salt concentrations.

To characterize the effects of ionic strength on fast timescale backbone dynamics of the bound peptide, we measured ¹⁵N backbone relaxation data. Heteronuclear ¹⁵N-{^1^H} NOE values, which are sensitive to picosecond timescale motions, showed little variation upon addition of salt (Figure 2B). Similarly, backbone amide order parameters (S²), which were derived from model-free analysis of the ^15^N relaxation data and report on the amplitude of motion on the sub-nanosecond timescale, were largely unaffected by increasing ionic strength (Figure 2C). The only salt-dependent effects observed are confined primarily to some of the peripheral lysine residues, while the core binding motif, the PxxP-containing region exhibits no significant change in heteronuclear NOE or S² values. This is in contrast to previous work showing that changes to the ArkA sequence have a large effect on the dynamics of the bound complex and the conformation of the AbpSH3 domain^10^. Although the NMR backbone dynamics data show that the peptide termini are more flexible than the central binding motif, this flexibility changes only slightly at increased salt concentration.

We also performed MD simulations of the ArkA-ApbSH3 complex to examine changes in conformation or dynamics under different salt conditions. We performed 10 independent simulations of both the ArkA17 and ArkA12 bound complexes domain under no salt and 800 mM NaCl conditions. Atomic fluctuations were used to quantify the dynamics of the protein backbone during the simulations. Overall, we saw only a slight difference in ArkA and AbpSH3 backbone fluctuations between the simulations at different salt concentrations, consistent with the order parameters from NMR (Figure 2D and Figure S4D). We also observed no large changes in the intermolecular contacts between ArkA and AbpSH3 in salt compared to no salt by NMR and MD (Figure S4E).

Attractive short and long-range electrostatic interactions between charged residues of AbpSH3 and ArkA were measured using MD simulations, which show a large network of contacts (Figure 1A). The central lysine, K(-3), is the only lysine that is involved in a stable short-range interaction (with E17), while the periphery lysines are predominantly involved in transient and long-range contacts (Figure 3). There are slightly fewer interactions with increased salt and a small shift from short-range to long-range, indicating that there are subtle changes in interactions even though the bound peptide structure and dynamics are generally unchanged by salt, consistent with the NMR data (Figure S5). This suggests that salt would not have a large effect on the ArkA dissociation rate. This prediction was experimentally tested in the following section with kinetics measurements at varying salt concentration.

**Fig 3:**
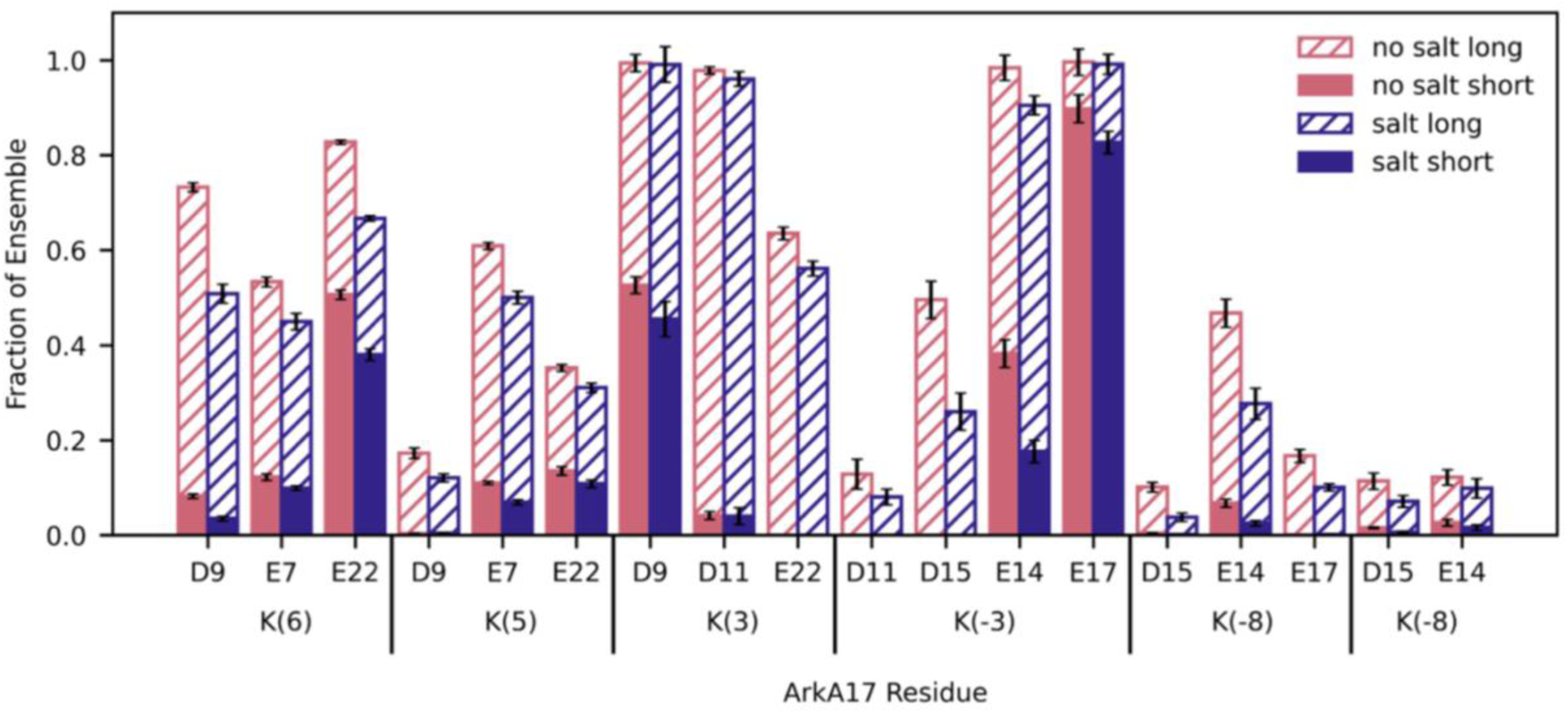
**The AbpSH3-ArkA complex contains a robust network of electrostatic interactions**. The populations of long (hatched) and short (solid) electrostatic interactions without (pink) and with (blue) salt from MD simulations of the bound ArkA17-AbpSH3 complex for each of the six lysine residues in the peptide. Each set of bars represents the population of a specific interaction between an ArkA lysine (bottom label) and a AbpSH3 aspartic acid or glutamic acid (top label).

### 3.3 Association rate is very fast due to long-range electrostatics

To characterize the overall transition state for binding (†), we measured the binding kinetics of the ArkA12 and ArkA17 peptides to AbpSH3 at different concentrations of NaCl using CPMG NMR with complexes that have approximately 5-10% peptide bound. Both peptides led to similar chemical shift perturbations in the domain at all salt concentrations confirming the same binding surface in all complexes and conditions (Figure S4A-C). The measured CPMG exchange rate constants and populations of AbpSH3 bound to peptide allowed us to calculate the association and dissociation rate constants (*k_on_* and *k_off_*) using our *K_D_* from ITC. At low ionic strength, we hypothesized that the transition state for ArkA binding to AbpSH3 would be stabilized by long-range electrostatic attraction and there would be fewer additional favorable electrostatic interactions formed after the transition state. Consistent with the results in Gerlach *et al*. 2020^17^, we found association rates at low salt of 1.21 x 10^9^ M^-1^ s^-1^ and 2.94 x 10^9^ M^-1^ s^-1^ for ArkA12 and ArkA17, respectively (Table 4 and Table S1). These association rates are faster than diffusion, consistent with electrostatic enhancement of binding. We also measured a clear trend in the natural log of *k_on_* and *k_off_* for the two peptides with ionic strength, where ln*k_off_* increases and ln*k_on_* decreases with the square root of ionic strength (Figure 4). However, as predicted, salt affects association more than dissociation, for example, for ArkA17, the change in binding affinity is due to an 87-fold decrease in the association rate with a much smaller increase (3.7-fold) in the dissociation rate (Table 4). Therefore, the major effect of salt is on *ΔG_N_*_→†_ rather than *ΔG_B_*_→†_. We define a salt phi-value for peptide association,

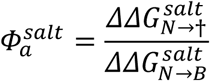

analogous to the salt phi-value for folding in our previous study of the effect of salt on AbpSH3 folding^7^. Where *ΔΔG^salt^* represents the difference in the barrier for association at 0 and 800 mM salt,

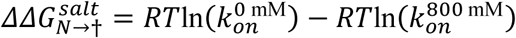

For ArkA17, 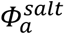 is 0.77 (Figure 5A and Table S1) and for ArkA12 it is 0.71, showing that the majority of the long-range electrostatic interactions that stabilize the AbpSH3-ArkA bound complex at low ionic strength are formed before the transition state, consistent with our hypothesis. Long-range electrostatic interactions are therefore clearly critical to AbpSH3-ArkA association and help to explain the very fast binding rate. This is consistent with previous studies that have found diffusion limited association of protein binding partners can be enhanced by long-range electrostatic interactions by a factor of 10^3^ or higher^3,5,32–35^. Combined with the free energy of AbpSH3 folding from our previous study^7^, we can see that the effect of salt on binding is caused by stabilization of the unbound AbpSH3 complex, *N* (Figure 5A). By lowering the free energy of the unbound domain, salt increases the barrier to binding ArkA and decreases the free energy benefit from reaching the bound state. The effect of salt concentration is different from that of point mutations that disrupt salt bridges, which have been shown to increase the dissociation rate for protein-protein interactions (low phi values)^33,36–39^, as these short-range interactions are geometrically constrained and form only in the fully bound state. Unlike point mutations, salt primarily screens long-range electrostatic interactions and therefore would not have a large effect on *ΔG_B_*_→†_ compared to *ΔG_N_*_→†_ for ArkA binding, as most of the long-range electrostatic interactions that ArkA forms with AbpSH3 are formed before the transition state. The favorable effect of salt on the free energy of the system is progressively smaller as ArkA-SH3 moves along the reaction coordinate toward the bound state, since salt has a smaller effect as ArkA interacts more with the SH3 domain (Figure 5A). Recent work by Álvarez *et al*. has used salt to deconvolute the effects of long-range electrostatic interactions and the remaining additional interactions for a charged residue mutation during a coupled folding and binding process^40^. Their study uses stopped flow fluorescence to show that many charged residues contribute stabilizing long-range electrostatic interactions in the transition state to binding but also participate in other interactions that are energetically frustrated in the transition state, an effect they term electrostatic compensation. Similarly, we observe that long-range electrostatic interactions between ArkA and AbpSH3 are one of the key interactions that help to stabilize the transition state (†) relative to the unbound state (*N*).

**Fig 4:**
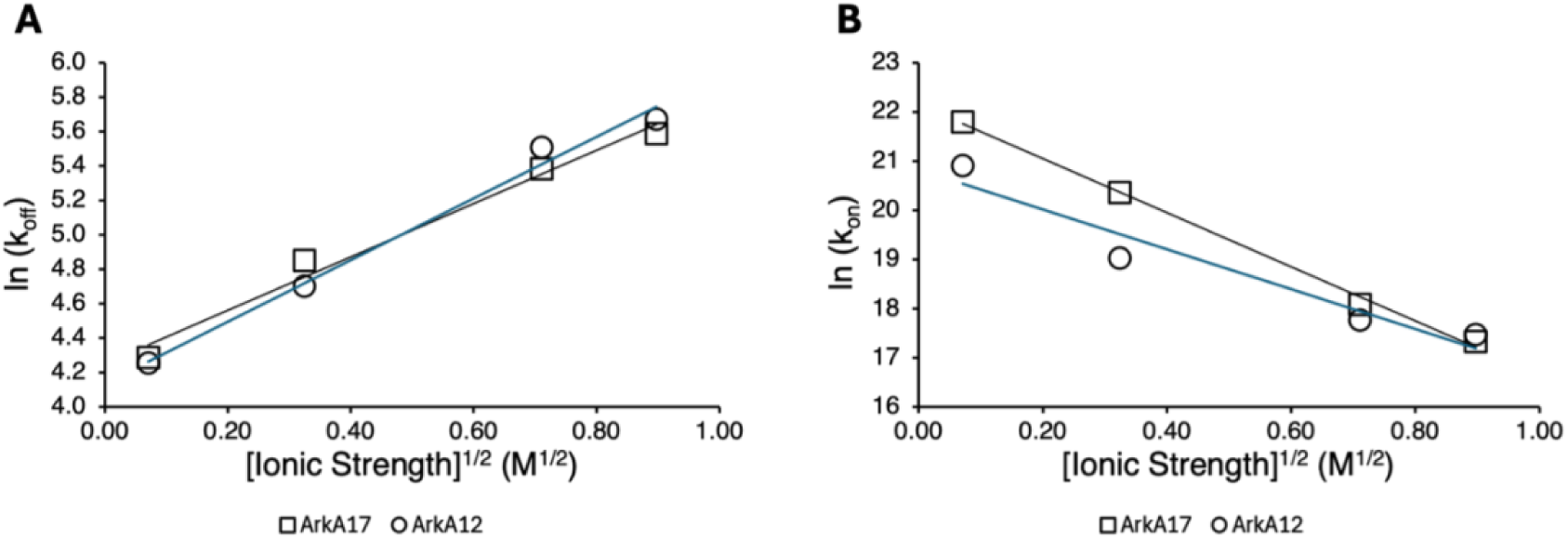
AbpSH3-ArkA binding kinetics at different ionic strengths. Natural log of (A) *k_off_* and (B) *k_on_* from CPMG experiments plotted versus the square root of ionic strength for ArkA17 and ArkA12. The y-axis range for the *k_on_* plot is larger, showing that association rate is affected more by ionic strength than dissociation rate.

**Table 4.** Experimental and simulation binding kinetics for ArkA12.

| | $K_D$ (μM) | $k_{off}$ (s<sup>-1</sup>) | | $k_{on}$ (s<sup>-1</sup>M<sup>-1</sup>) |
| --- | --- | --- | --- | --- |
| Exp. (0 mM NaCl) | 0.0582 | 70.3 | | $1.21 \times 10^9$ |
| Exp. (800 mM NaCl) | 7.51 | 290 | | $3.86 \times 10^7$ |
| | $K_D^{[EC]}$ (mM) | $k_1$<br>(10<sup>9</sup> s<sup>-1</sup>M<sup>-1</sup>) | $k_{-1}$<br>(10<sup>6</sup>s<sup>-1</sup>) | $k_{on}$<br>(10<sup>7</sup> s<sup>-1</sup>M<sup>-1</sup>) |
| MD (0 mM NaCl) | 0.50 ± 0.11 | 5.08 ± 0.72 | 2.54 ± 0.42 | 5.3 ± 1.8 |
| MD (800 mM NaCl) | 3.62 ± 0.75 | 1.41 ± 0.20 | 5.10 ± 0.78 | 5.3 ± 1.8 |

**Fig 5:**
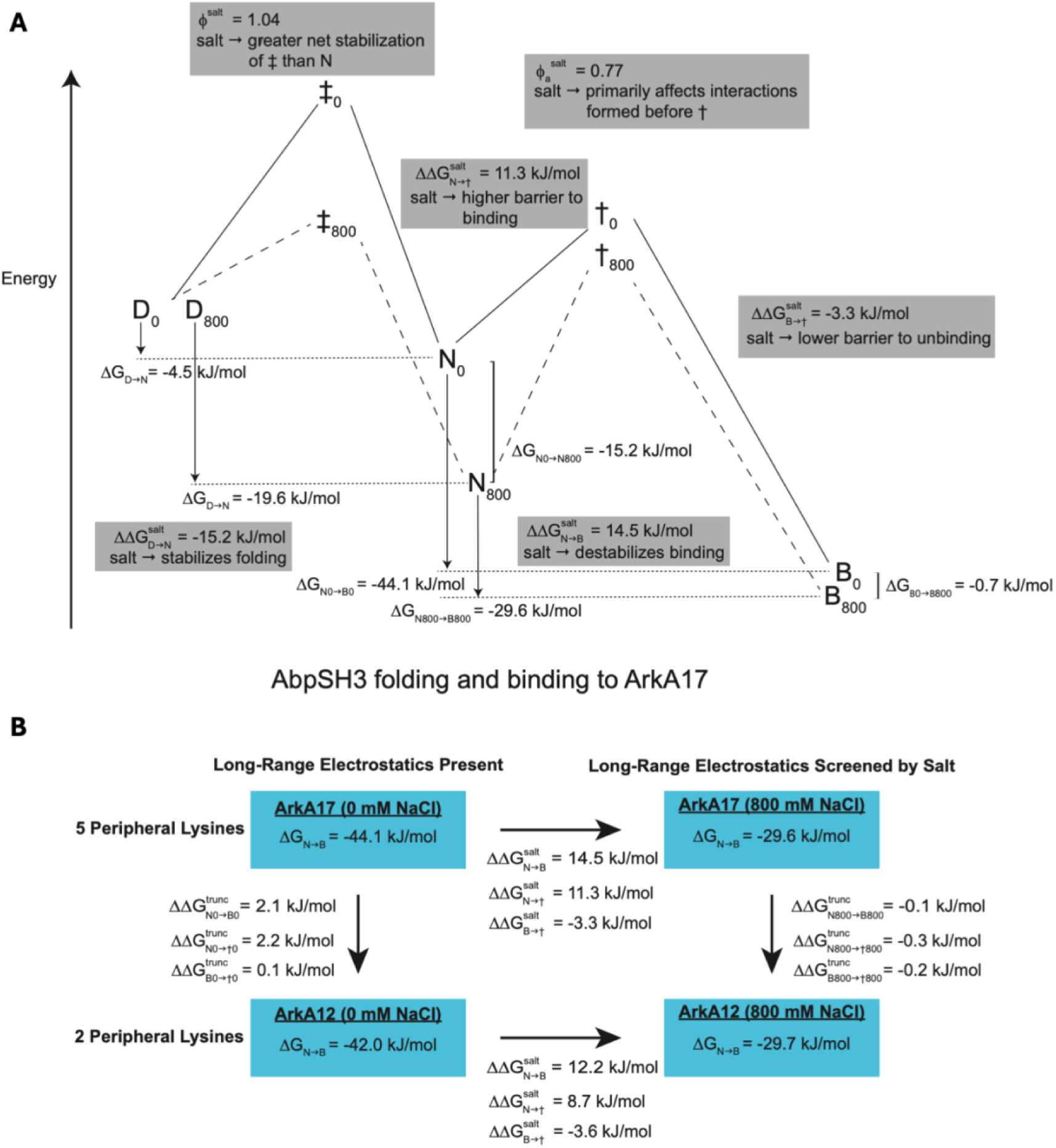
Summary of the role of electrostatics on ArkA-AbpSH3 binding kinetics and thermodynamics. (A) Free energy diagram of the folding of AbpSH3^7^ and binding to ArkA17 based on thermodynamic data from ITC and kinetic data from NMR CPMG experiments. *D* represents the denatured state of the unbound SH3 domain, *N* represents the native folded state of the unbound SH3 domain, *B* represents the bound complex, ‡ represents the transition state for folding, and † represents the transition state for binding. The subscripts 0 and 800 indicate the NaCl concentration. (B) Thermodynamic cycle based on experimental data for ArkA17 and ArkA12 binding at high and low salt. The superscript *salt* indicates a change in *ΔG* between 0 and 800 mM salt concentration, while the superscript *trunc* indicates a change in *ΔG* between ArkA17 and ArkA12 binding.

Comparing the kinetics of binding for ArkA17 and ArkA12, we are also able to learn about the role of the peripheral lysine residues in the binding process, as ArkA12 lacks three peripheral lysine residues and therefore contains a lower net charge (+3) than ArkA17 (+6). This leads to a lower association rate for ArkA12 at low salt concentrations (2.4-fold decrease). This difference disappears at high salt concentrations, where the effect of these additional lysines is no longer significant due to screening of long-range electrostatic interactions. Examining the *k_off_* values for the two peptides, we see no difference across all salt concentrations, indicating that the peripheral lysine residues do not play a significant role in dissociation. This is consistent with the bound state being stabilized by specific, short-range interactions in the core of the binding motif. Our simulation results support this picture, as they show that the only short-range electrostatic interaction that is formed consistently in the bound state is between E17 and K(-3), which is present in both peptides (Figure 3), and the core contacts in the binding motif are consistent for simulations of both the ArkA12 and ArkA17 complexes. Electrostatic interactions with the other lysine residues are transient and highly dynamic (Figure 1), consistent with the idea that fuzzy interactions (those interactions that are transiently occupied and switch between different interaction partners while the overall complex remains bound) involving peripherally charged residues can help to stabilize the SH3 bound state^41^. While these residues are critical to stabilizing binding, they are also present in the transition state. Therefore, removing those peripheral lysines in ArkA12 does not affect *ΔG_B_*_→†_ or the dissociation rate (Figure 5B). The change in *ΔG_N_*_→*B*_ going from ArkA17 to ArkA12 at low-salt is entirely due to a change in *ΔG_N_*_→†_, while at high salt there is no change in *ΔG_N_*_→*B*_ going from ArkA17 to ArkA12, as the effect of the peripheral lysines on the association rate are effectively screened by salt. While we do not have kinetics data for the ArkA10GS construct, from the ITC data we can see that removing all five peripheral lysines has an even more dramatic effect on binding, but this effect is entirely screened by salt (Figure 1B), and likely also primarily affects the associate rate. We propose that this is a commonality among peripheral charged residues in the binding of disordered sequences. While the long-range fuzzy electrostatic interactions formed by these residues can be important during the initial stages of binding, they should already be formed in the transition state. These peripheral charged residues can help to stabilize the transition state, but since they do not form geometrically constrained interactions in the bound state, we do not expect them to engage in new non-electrostatic interactions that lead to electrostatic compensation of frustrated transition state energetics as seen by Álvarez *et al*., who observed this effect for charged residues in the more structured core of the motif^40^. Our simulations of the binding pathway, including the encounter complex intermediate, allow us to further examine where electrostatic interactions form during association and how this is affected by salt.

### 3.4 Encounter complex stability and interactions are sensitive to salt

SH3 domain binding occurs via an encounter complex intermediate (eq 2), and examining the effect of salt on the encounter complex in our MD simulations can help us understand the role of electrostatic interactions during association in more detail. Simulations of ArkA12 binding in the absence of salt previously showed that the rates of encounter complex formation, *k*_1_, and encounter complex dissociation, *k*_−1_, are orders of magnitude faster than the transition to the fully bound state, *k*_2_ ^17^. This allows us to apply the pre-equilibrium approximation to describe the two-step binding, as the unbound state and encounter complex interconvert rapidly forming a pre-equilibrium before the encounter complex progresses to the bound state. The dissociation rate, *k_off_*, is still orders of magnitude slower, allowing formation of the bound state to be approximated as an irreversible step:

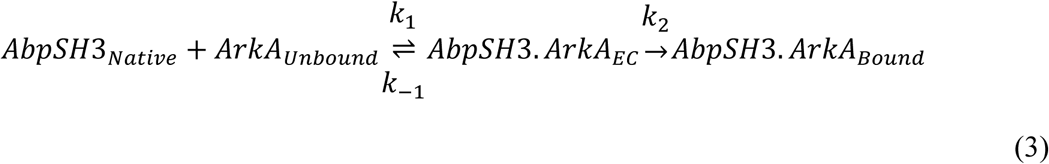

where *k*_1_ >> *k*_2_ and *k*_−1_ >> *k*_2_. Using this approximation,

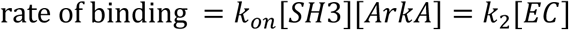

and

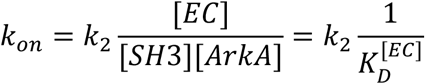

where 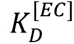 is the pseudo dissociation constant for the encounter complex. Therefore, the overall rate constant, *k_on_*, depends on *k*_1_, *k*_−1_, and *k*_2_ as in

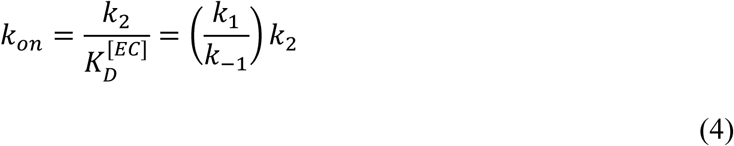

Since the simulations of ArkA12 binding in the absence of salt showed that long-range electrostatic interactions form during step 1 and are already present in the encounter complex^17^, we expected salt would primarily affect the first step of encounter complex formation in our simulations. We performed 50 independent binding simulations of ArkA12 and AbpSH3 in 800 mM NaCl and compared them to our simulations in the absence of salt and found that there is strong evidence that *k*_1_ decreases in the presence of salt based on the Bayesian odds ratio (Table S3). Additionally, *k*_−1_ increases with high salt (Table 4), leading to a lower population of the encounter complex in the pre-equilibrium that is formed before step 2 (Table 4). Similarly to the bound state, the encounter complex binding free energy, *ΔG_N_*_→*EC*_ decreases in magnitude in the presence of salt because the unbound SH3 domain is stabilized more by salt than the encounter complex is, increasing 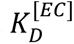 . A destabilized encounter complex will contribute to the overall lower association rate that we observe experimentally in the presence of salt, since *k_on_* has an inverse relationship with 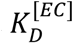 (Eqn. 4). Surprisingly, we do not observe a corresponding decrease in *k _on_*in our MD simulations (Table 4), which we estimate from the small number of simulations (9 out of 50) that complete the entire binding process. However, this number of binding events is underpowered to determine if there is indeed a difference in the association rate caused by salt (odds ratio of 0.38)^30^. Therefore, we focus on the effect of salt on encounter complex formation and its ensemble when interpreting our binding simulation data.

Structural analysis from our simulations can reveal how the encounter complex conformational ensemble is affected by salt. The difference contact map in Figure 6A highlights the effect of salt to both native and nonnative interactions between ArkA and the AbpSH3 in the encounter complex. In general, the encounter complex forms transient and non-specific interactions both with and without salt present and, like for the bound complex, there is only a slight decrease in the average number of contacts with salt (Figure 6A and Table S2). However, when comparing the total number of electrostatic interactions that are disrupted by salt, we see that the encounter complex is more impacted than the bound state (Figure 6B and Table S5). With salt present, long-range electrostatic interactions formed between the peptide and domain are not as thermodynamically beneficial because salt ions can form equivalent types of interactions with the charged proteins. This is true in both the bound state and encounter complex, but in the bound state the long-range interactions between ArkA and the SH3 domain remain even in the presence of salt because the core motif has locked onto the binding surface and the peptide is geometrically constrained. In the encounter complex, there is no such geometric constraint (the salt-bridge between K(-3) and E17 is rarely formed^17^) and therefore the presence of salt leads to more sampling of states that do not involve these long-range electrostatic interactions between ArkA and the SH3 domain, changing the conformational ensemble of the encounter complex (Figure S6).

**Fig 6:**
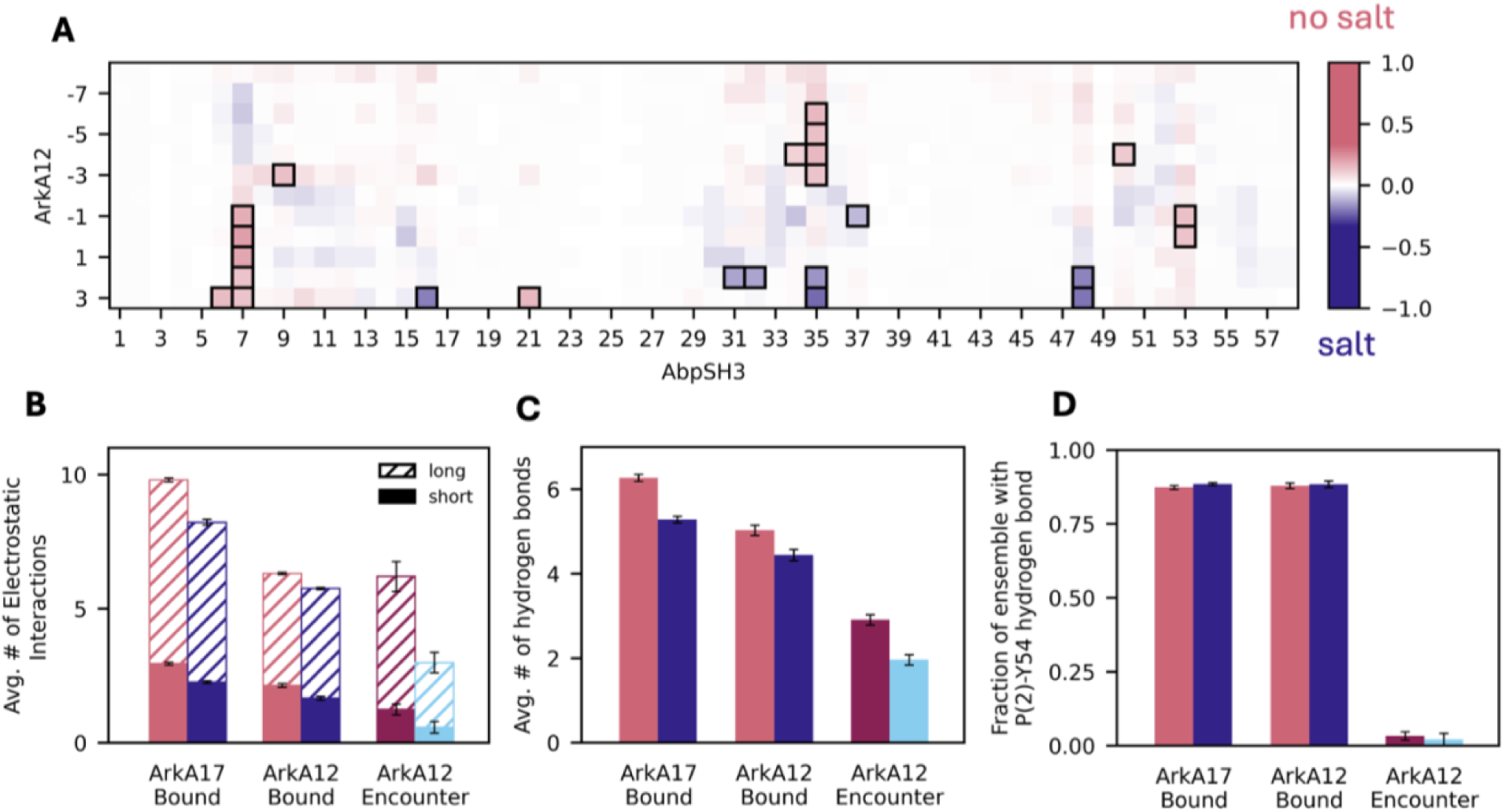
Salt disrupts the encounter complex more than the bound state. (A) Contact difference map from simulations of the ArkA12 encounter complex with and without salt. Pink contacts are occupied more frequently in the simulations without salt and blue contacts are occupied more frequently in simulations with 800-mM NaCl present. Boxed contacts indicate differences that have a p-value below 0.05. (B) Average number of electrostatic interactions (solid bars for short-range interactions, hashed for long-range interactions) in the simulated bound state (ArkA17 and ArkA12) and encounter complex (ArkA12) with (pink or red bars on left) and without salt (blue bars on right). (C) Average number of h-bonds in the bound and encounter complex ArkA12 MD simulations with (pink or red bars on left) and without salt (blue bars on right). (D) P(2) to Y54 h-bond in bound and encounter complex ArkA12 MD simulations with and without salt. P-values for differences between no salt and salt for panels B and C are given in Table S5 and Table S6.

In addition to interactions between charged residues, ArkA and AbpSH3 can form hydrogen bonds, which are also electrostatically based short-range interactions. Our MD simulations show a small decrease in the average number of hydrogen bonds for both the encounter and bound complexes when salt is present (Figure 6C and Table S6). However, no effect of salt was observed on the stable P(2) to Y54 hydrogen bond that is formed in step 2 and is present in over 80% of the bound state ensemble (Figure 6D), showing that salt only affects the hydrogen bonds that are in more dynamic exchange. Likewise, hydrophobic interactions are not affected by salt at any stage of the binding pathway, resulting in the same amount of buried surface area and the same hydrophobic interactions in the complexes with and without salt present (Figure 7 and Table S7).

**Fig 7:**
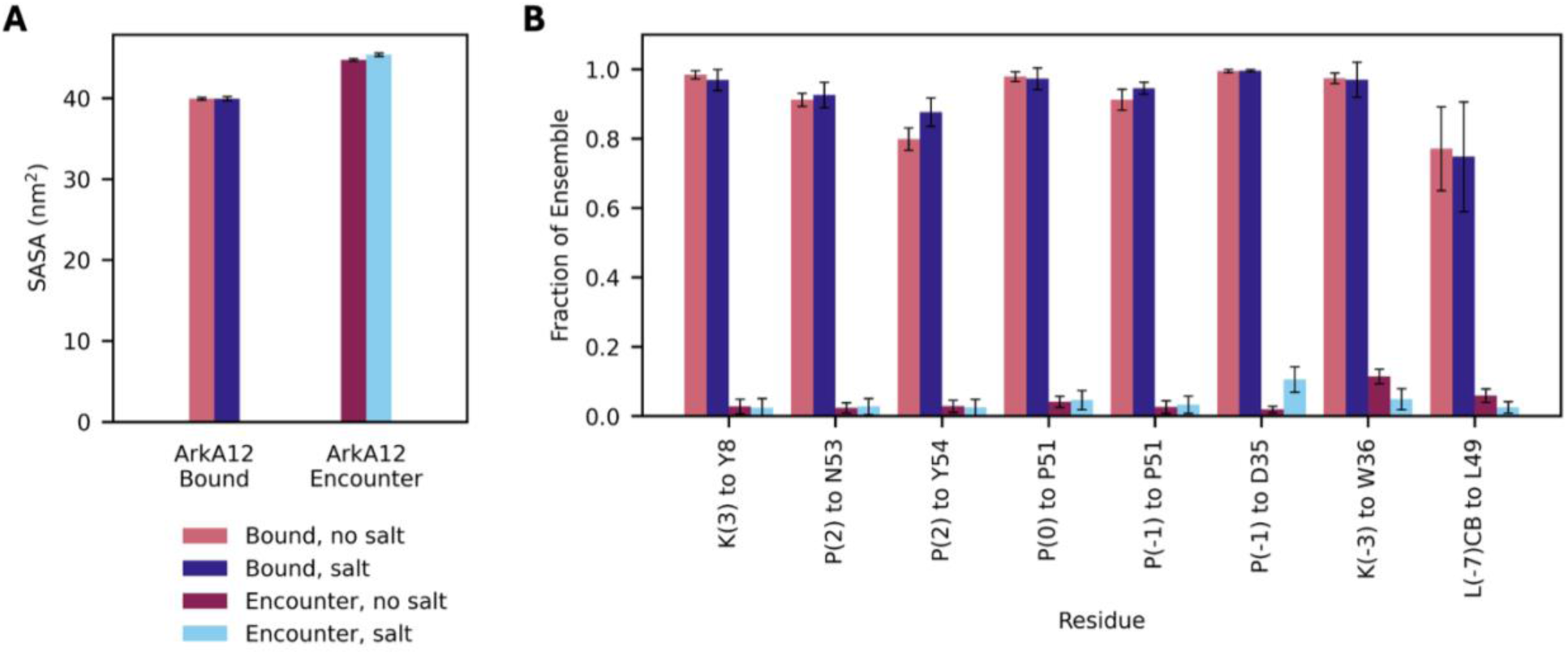
Hydrophobic interactions are not affected by salt. (A) Solvent accessible surface area from MD simulations of ArkA12 bound and in the encounter complex with salt (pink and red bars) and without salt (blue bars). (B) Occupancy of specific hydrophobic contacts present in the ArkA17-AbpSH3 complex NMR structure from MD simulations of ArkA12 bound and in the encounter complex with salt (pink and red bars) and without salt (blue bars). P-values for differences between no salt and salt for panel A are given in Table S7.

Overall, salt has a greater effect on the long-range electrostatic interactions formed in the encounter complex than on the electrostatic interactions formed in the bound state. These encounter complex electrostatics impact 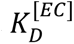 (and possibly *k_2_*) and therefore the association rate (eq 4), consistent with the CPMG NMR data. Ionic strength also influences the dissociation rate because salt stabilizes the transition state more than the bound state of the complex, although this effect is small, as reflected in the 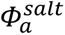 value of 0.77 (Figure 5A). While most of the long-range electrostatic interactions in the complex are present in the encounter complex and in the transition state, the salt dependence of the dissociation rate shows that there are some additional electrostatic interactions (perhaps hydrogen bonds or short-range electrostatics) present in the bound state that are not formed in the transition state. When salt ions are present to substitute for these interactions, there is a smaller energy penalty for disrupting them and the complex can more easily move from the bound state to the transition state.

The results of our binding simulations in the presence of salt are consistent with our previous conclusions from Gerlach *et al*. 2020 that long-range electrostatic interactions form primarily in step 1, while shorter-range interactions contribute more in step 2^17^. Both types of interactions have critical contributions to the ArkA binding affinity. Decreasing the number of peripheral lysine residues and adding salt both significantly decrease the binding affinity, demonstrating the importance of long-range and fuzzy interactions, which are formed in step 1. This supports the conclusions of Shen et al. that fuzzy, long-range electrostatic interactions are critical to the high affinity binding of the PRM NS1 to its SH3 binding partner^41^. It is important to note, however, that short-range interactions, formed in step 2, are also important for contributing to binding affinity as they stabilize the bound state compared to the encounter complex. This is supported by a previous study that mutated the central K(-3) residue, disrupting the dominant short-range electrostatic interaction and dramatically decreasing binding affinity^9^. The formation of a heterogenous ensemble with fuzzy, long-range electrostatic interactions in step 1, followed by short-range, geometrically constrained interactions forming in step 2 is likely a common feature of IDP binding pathways. Salt and peripheral charge mutations primarily affect the association rate because they impact the long-range electrostatic interactions that form in step 1. The encounter complex ensemble is heterogeneous and not geometrically constrained, which allows binding to occur more rapidly by lowering the entropic barrier for association. Non-electrostatic point mutations would not be expected to destabilize the encounter complex because it is a fuzzy complex with no specific interactions. While adding salt does destabilize the encounter complex, the ensemble nature of this intermediate also means that it can shift its ensemble to buffer against perturbations, dampening this effect and retaining a fast association rate. As a result, we see a smaller electrostatic enhancement effect for ArkA-AbpSH3 binding than that which is observed for the binding of similarly fast-binding highly-charged globular proteins that do not pass through a heterogeneous encounter complex intermediate to reach the bound state^3,5,34,35,42^. In ArkA binding, the encounter complex forms rapidly even without electrostatic enhancement, due to the absence of a large entropic penalty, so the electrostatic attraction between the partners can only increase the association rate by a smaller factor. Functionally, this rapid encounter complex formation could allow SH3 binding to proceed quickly in many different environments and cellular conditions, including different solute concentrations, post-translational modifications, and mutations; resulting in a more resilient fast association than the interaction between two folded domains, but a lower binding affinity.

### 3.5 Salt disrupts binding primarily through an enthalpic effect

Our ITC data at different salt concentrations combined with MD simulations that explicitly show ion interactions with the protein allow us to examine the mechanism by which salt affects ArkA-AbpSH3 binding in more detail. As discussed in section 2.1, salt stabilizes the unbound state of AbpSH3 more than it stabilizes the ArkA-AbpSH3 complex, which results in a decreased binding affinity with increased salt concentration (Figure 5). Figure 1C shows that this effect is the result of a change in enthalpy, with the enthalpy of binding becoming less favorable at higher salt concentrations. Mechanistically, this can be explained by the reduced number of favorable electrostatic interactions in the unbound versus bound states. At low salt concentrations, AbpSH3 does not form many favorable electrostatic interactions in the unbound state, but forms more favorable short and long-range electrostatic interactions when ArkA is bound, leading to a favorable enthalpy change. At high salt concentrations, however, this effect is less pronounced since salt ions are able to form favorable electrostatic interactions with AbpSH3 and ArkA even in the unbound state. Binding continues to be overall enthalpically favorable even at 800 mM NaCl because of stable short-range interactions that can form between the proteins but not with the salt ions. Changes in water enthalpy on binding could also potentially affect binding enthalpy.

The effect of salt on the entropy of binding is less straightforward for the ArkA-AbpSH3 system. Entropy of binding becomes more favorable as salt concentration increases, partially counteracting the unfavorable effect of salt on the enthalpy of binding (Figure 1D). This indicates that ion release is not a major contributor to the overall entropy of binding. Previous studies of DNA-protein binding and of interactions between two highly charged IDPs have shown that as bound salt ions are released from the protein or DNA during protein binding, this leads to a large increase in the entropy of those ions, which is a major contributor to the overall binding affinity^15,43^. In those cases, increasing salt concentration leads to a decrease in the favorable entropy of binding, which leads to a lower binding affinity at high salt. This is a result of the smaller contribution that each individual ion being released has to the overall entropy if there are a higher number of ions present in the solution. In the AbpSH3 system we see an opposite trend in binding entropy with ionic strength, which has also been seen in other studies^44,45^, indicating that ArkA-AbpSH3 binding does not result in a large release of ions.

We examined our high-salt MD simulations to better understand ion release with peptide binding from an atomistic perspective. Figure 8 shows that cations interact with many of the acidic AbpSH3 residues in both the unbound, encounter complex, and bound simulations. While the different residues on the surface vary in how frequently they form an ion contact, for most residues the time spent interacting with salt ions does not change much with ArkA binding, except for E14 and E17. This is consistent with our observation that most electrostatic interactions between ArkA and AbpSH3 are long-range and transient (Figure 3 and Figure 6), providing an opportunity for ions to also transiently interact with the charged residues. In the case of E14 and E17, which have a cation bound 45% and 26% of the time respectively in the apo state, we observe a decrease to 28% and 5% when the ArkA K(-3) residue forms a salt bridge at that location in the bound state. Because the AbpSH3 domain is so highly charged and contains 15 acidic residues on its surface, the decrease in interaction with salt ions at E14 and E17 with ArkA binding is a small change relative to the total number of ions interacting. Each acidic residue also interacts with a high number of different cations over the course of the simulation, indicating that the ions are exchanging rather than remaining statically bound (Table S8). Overall, AbpSH3 ion interactions engage in non-specific territorial ion binding that results in a loosely associated net positive ion atmosphere in the 3 Å surrounding the complex, which ranges from 9.8 in the apo state to 7.9 in the presence of ArkA17 (Table 5).

**Fig 8:**
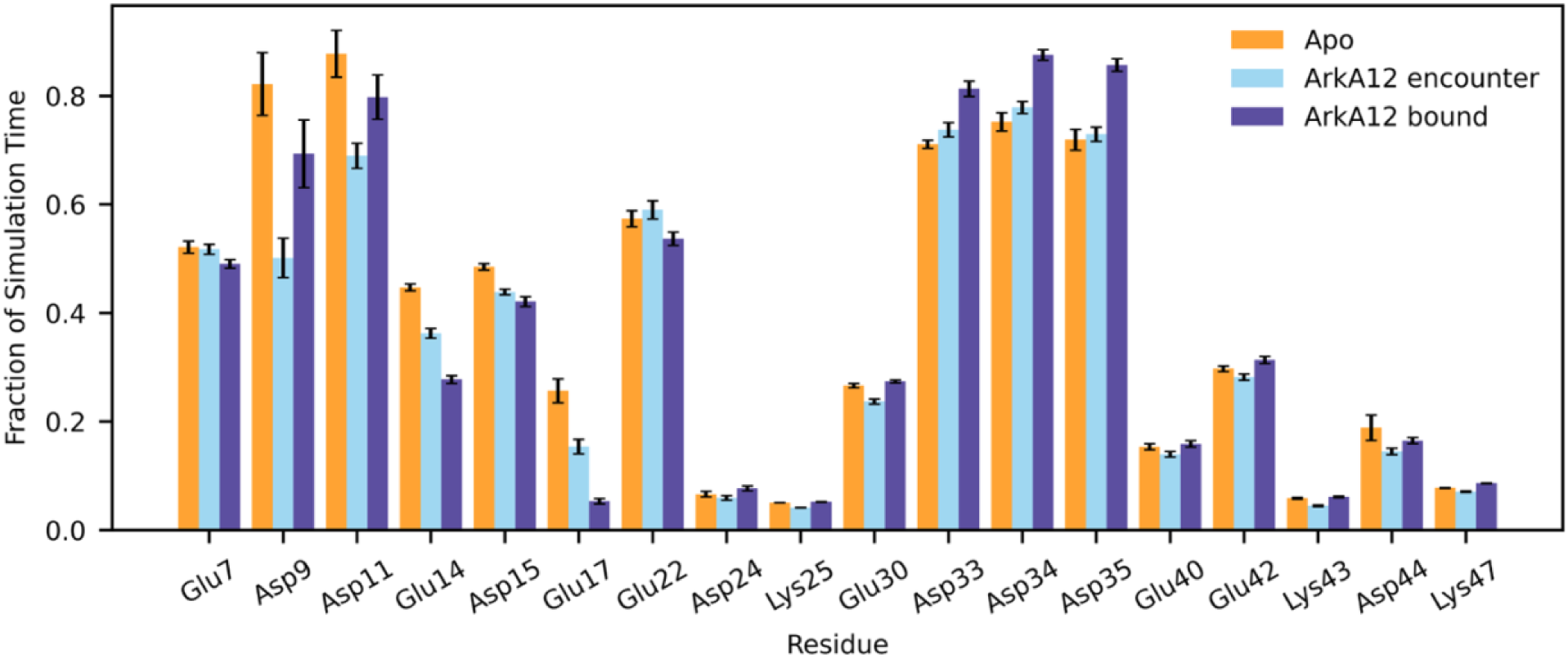
Ion contacts by residue. Fraction of the simulation time that the oppositely charged ion (Na^+^ for acidic residues and Cl^-^for basic residues) spends in contact with the charged group of the SH3 domain residues in high salt simulations. Ion contact time is shown for the simulations of the apo AbpSH3 domain (orange), ArkA12 encounter complex (light blue), and ArkA12 bound state (dark purple).

**Table 5.**
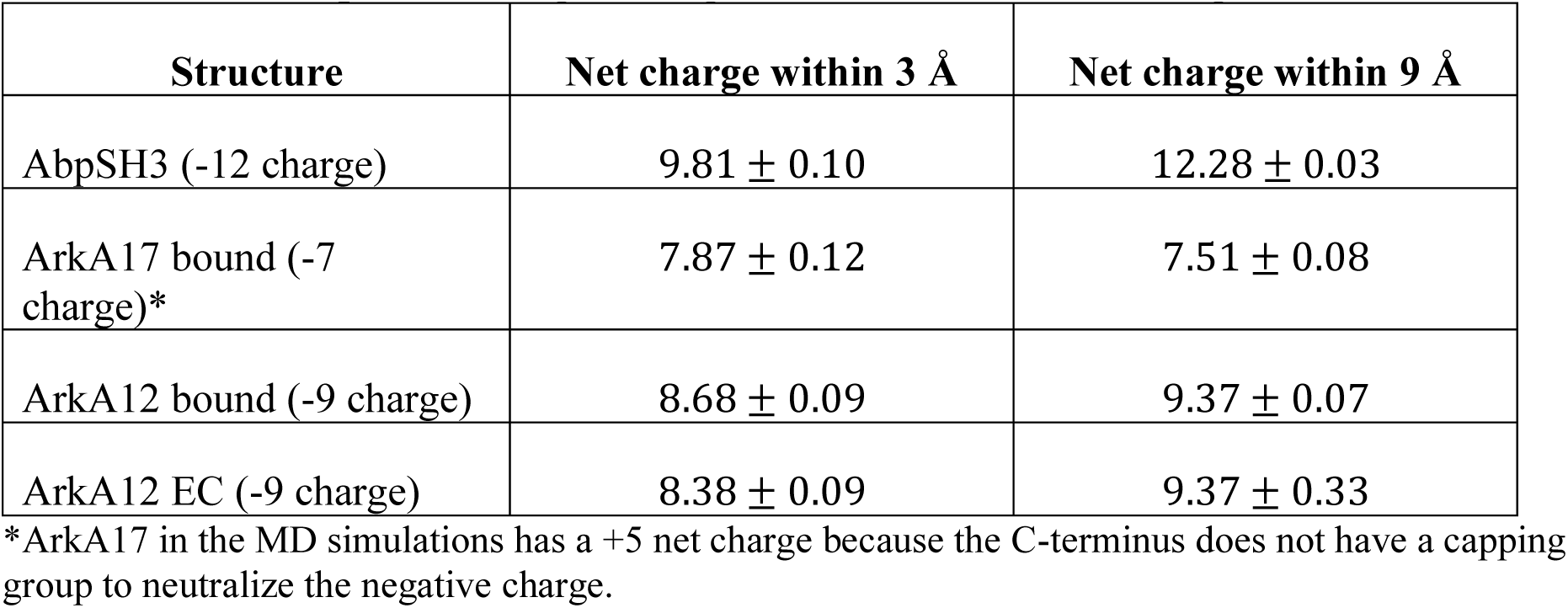
Ion atmosphere during binding from MD simulations in high salt. ArkA17 in the MD simulations has a +5 net charge because the C-terminus does not have a capping group to neutralize the negative charge.

| Structure | Net charge within 3 Å | Net charge within 9 Å |
| --- | --- | --- |
| AbpSH3 (-12 charge) | $9.81 \pm 0.10$ | $12.28 \pm 0.03$ |
| ArKA17 bound (-7 charge)* | $7.87 \pm 0.12$ | $7.51 \pm 0.08$ |
| ArKA12 bound (-9 charge) | $8.68 \pm 0.09$ | $9.37 \pm 0.07$ |
| ArKA12 EC (-9 charge) | $8.38 \pm 0.09$ | $9.37 \pm 0.33$ |

Taken together, our data do not support the ion release model of tightly bound, low-entropy ions that completely dissociate into bulk solvent on binding, leading to a large entropy increase. Instead, they support a model of loosely associated, exchanging ions in the unbound state and a small or insignificant increase in ion entropy on binding, as the ion atmosphere loses just a few cations. The favorable increase in entropy of binding observed with increased salt concentration may be a result of changes in water entropy, which we have not measured directly. Protein conformational entropy could also contribute to this change. Our MD simulations and NMR data do not show evidence for a large change in conformational entropy of the domain or peptide in the bound state with salt concentration. However, salt could affect the entropy of the unbound ArkA peptide, which is consistent with a study by Vancraenenbroeck et al. that shows how salt affects binding affinity by influencing the unbound IDP conformation^16^. We have not examined the effect of salt on the unbound ArkA in our simulations; however, if increased salt helps to structure ArkA and reduce the entropy of the unbound ArkA, this would result in a favorable reduction in the loss of conformational entropy of the peptide on binding, explaining the trend in *ΔS* of binding with ionic strength (Figure 1D). Ultimately, the primary effect of salt on ArkA-AbpSH3 binding is by disrupting enthalpic interactions, with a smaller overall effect on the binding entropy.

## 4. Conclusion

Using salt concentration as a probe, we have examined the role of long-range electrostatic interactions in the binding pathway of the highly acidic AbpSH3 domain. Although salt has been previously used to investigate electrostatic effects for protein-protein interactions, there are still many open questions about how these effects operate in the context of binding to a disordered sequence, such as the AbpSH3 intrinsically disordered partner ArkA. Through our ITC and NMR experiments and MD simulations, we have shown that the AbpSH3-ArkA binding process has several differences from previously studied interactions between two highly-charged folded domains.

Firstly, we are able to reveal why the **electrostatic enhancement** of the binding rate for SH3 domain-peptide interactions is less pronounced than in the case of similarly fast-binding *folded* charged proteins. For the AbpSH3-ArkA system, binding occurs in two steps by way of an encounter complex intermediate that contains many transient long-range electrostatic interactions. Salt destabilizes this intermediate by substituting for the long-range charged interactions made between the peptide and the domain. Therefore, in the presence of high salt, the encounter complex forms more slowly and dissociates more easily. However, the encounter complex is highly heterogenous and is still able to form in the presence of salt due to the low entropic penalty for binding. Once the encounter complex forms, the transition to the bound state is facilitated by the proximity of the peptide and binding surface. Therefore, the heterogeneous encounter complex serves as a buffer to perturbations, allowing resilient fast association that relies less on electrostatics than for folded charged domains, but with a lower binding affinity.

Secondly, although the ArkA peptide contains several positively charged residues that interact favorably with the domain through electrostatic interactions, these residues play different roles in the binding process. Experiments with different ArkA constructs in high salt show that the **peripheral lysine residues** are only involved in long-range electrostatic interactions that are disrupted by salt. These long-range electrostatic interactions form during the first step of binding, limiting the effect of the peripheral lysines to the association rate. Only the central K(-3) ArkA residue forms short-range electrostatic interactions that are not present in the encounter complex ensemble. Peripheral lysines therefore present an effective way for evolution to modulate the binding pathway without changing dissociation. When two folded domains form a complex, the electrostatic interactions are generally specific salt bridges that form at close range and likely explain the higher binding affinity but also affect the dissociation rate. In an IDP recognition interaction, we see that charged residues peripheral to the core binding motif have a more limited impact, affecting the binding affinity only through the association rate by forming fuzzy interactions in the encounter complex and bound state.

Thirdly, the dynamic and transient nature of the majority of the electrostatic interactions formed on a relatively small ArkA-AbpSH3 binding interface means that **ion release** does not significantly contribute to a favorable entropy of binding. For highly charged proteins that bind to DNA, ion release upon binding is a major contribution to the favorable thermodynamics of binding due to the increase in entropy of the released salt ions. In the case of AbpSH3 binding to ArkA, despite the high overall charge of the SH3 domain, ion release does not significantly enhance binding affinity. Only one of the ArkA lysine residues forms a stable salt bridge with the SH3 domain, displacing an Na^+^ ion, while the other electrostatic interactions are transient and fluctuating. The resulting effect is a relatively small decrease in ion interactions upon ArkA binding, and overall the binding of the complex is entropically unfavorable. The high net charge of the SH3 domain contributes to a favorable binding affinity that decreases with salt concentration, but this is an enthalpic rather than entropic effect.

In our previous paper on the folding of the highly charged AbpSH3 domain, we hypothesized that the conserved high negative charge of the AbpSH3 domain is selected for because of its ability to enhance the association rate with the disordered binding partner^7^. Now we can add nuance to that picture; the high charge does not enhance association as much as it would for two folded proteins binding, but the combination of a disordered binding partner and many fuzzy, long-range electrostatic interactions has other benefits. In the context of the complex cellular environment, electrostatic steering would help the domain and disordered binding partner come into closer proximity. The AbpSH3 net dipole would steer a positively charged disordered region to interact preferentially with the binding surface, and the encounter complex would form without a large entropic barrier. The presence of many non-specific, fuzzy interactions, both electrostatic and hydrophobic, would allow the domain and IDR to remain in an encounter complex that is rapidly sampling different regions until the target motif exchanges onto the binding surface, rather than diffusing away from each other (Figure 9). Overall, the role of electrostatics is similar to the dimensionality reduction from a 3D to 1D search in transcription factor binding to DNA^40,46^. In our proposed mechanism, a heterogeneous encounter complex forms quickly through electrostatic steering and allows the SH3 domain to interact with a range of positively charged IDRs, conferring a functional advantage for quickly finding its unique target sequence *in vivo*.

**Fig 9:**
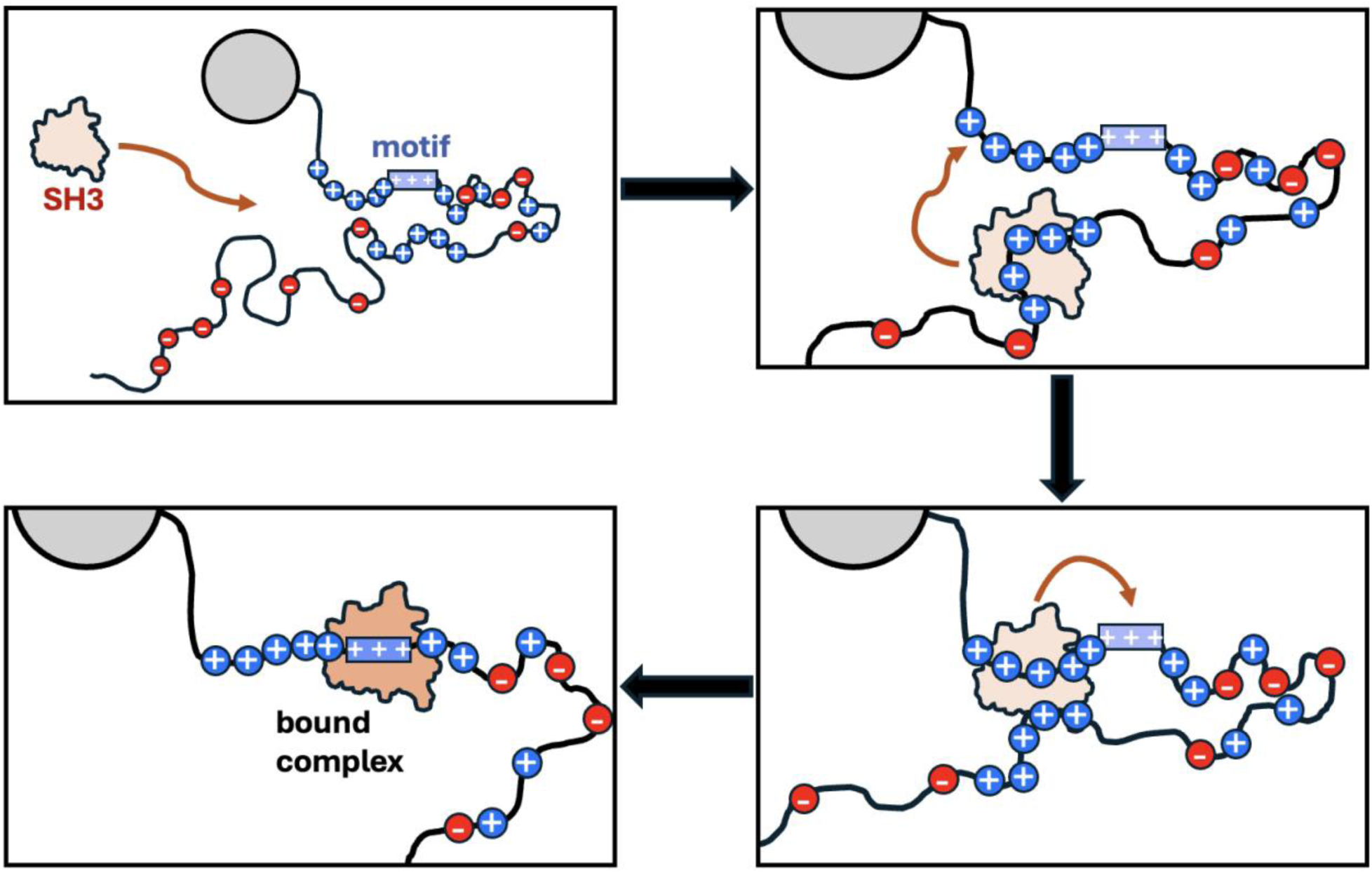
Cartoon of how a highly-charged SH3 domain could search for its target domain in the cellular environment. The SH3 binding motif (ArkA in this study) is represented with a rectangle in a long disordered region of the SH3 partner protein. Panel 1 is the initial unbound state. Panel 2 is the formation of a heterogeneous encounter complex ensemble with a positively charged disordered region that does not contain the motif. In panel 3, the SH3 domain diffuses to a different positively charged region while remaining in the encounter complex. In panel 4, the SH3 domain reaches the partner motif and locks into the thermodynamically stable bound state, while still engaging in fuzzy interactions with peripheral lysines.

## Supporting Information Description

Additional data from ITC, NMR, and MD simulations

Complete data table from all ITC experiments

All chemical shift values for the AbpSH3 domain from peptide-saturated complex for ArkA17, ArkA12, and ArkA10GS, each at 0 mM, 0.1 mM, 0.5 mM, and 0.8 mM NaCl

## Supporting information

ITC data table

AbpSH3 domain chemical shifts data table

## Acknowledgement

We would like to thank Nick Razo and Sam Hawke for consultation on statistical analysis of MD simulations. Thank you to Sophia Liras for assisting with protein expression and purification. Thank you to Michael Donnelly for computational support. K.A.B. thanks the MERCURY Consortium, supported by National Science Foundation grant CHE-2320718, for mentoring support. This work was supported by National Science Foundation grant MCB-2324974 to K.A.B. and National Institute of General Medical Sciences grant R35GM128906 to M.P.L. Support was provided by the Schupf Scholar Program to A.M., D.P., and S.R. and by the Skidmore College Faculty Student Summer Research Program to O.O. and C.M.

