## Supporting Information for "How a highly acidic SH3 domain binds to its intrinsically disordered partner through the formation of an encounter complex intermediate"

**Text S1:** Details of $^{15}N$-labelled AbpSH3 domain expression and purification

For NMR CPMG and HSQC experiments, the $^{15}N$-labelled AbpSH3 domain was expressed in minimal media (supplemented with $^{15}N$-labelled ammonium chloride). The same plasmid that was used for the unlabeled domain was transformed into NiCo21 (DE3) competent E. Coli cells for 15N-labeled expression. A 60-mL starter culture was grown in Terrific Broth media using 1-2 colonies from the transformation plate. After growing for 6 h at 37 °C at 225 rpm, the starter culture was centrifuged to remove the media (1800 rpm for 10 minutes). and transferred to a 200-mL minimal media culture based on the M9 minimal media, but with increased nutrient concentration (113 mM $Na_2HPO_4$, 69 mM $KH_2$ $PO_4$, 18.8 mM NaCl, 2.7 mM $CaCl_2$, 4 mM $MgSO_4$, 22.2 mM glucose, 81.9 mM biotin, 1.02 g/L $^{15}NH_4Cl$, 2x MEM vitamin solution, and 1x trace metals solution)[18]. After overnight growth at 37 °C at 225 rpm, this culture was centrifuged to remove the media (1800 rpm for 20 minutes) and split between four large flasks containing 450 mL of minimal media. The cells were allowed to grow at 37 °C at 225 rpm until reaching an OD between 1.8 and 2.5. Once the target OD was reached, IPTG was added to a concentration of 1 mM to induce the cells and express the AbpSH3 protein. After induction the cells were grown at 37 °C at 225 rpm for 4 hours and then harvested through centrifugation.

The $^{15}N$-labelled AbpSH3 domain was purified using a denaturing purification[17]. The cell pellet suspended in 2.5 mL/g of cold lysis buffer (10 mM $Na_2HPO4$, 10 mM Tris, 6.0 M of GuHCl, and 10 mM imidazole, pH 8). The cells and buffer were vortexed to homogeneity; they were then sonicated twice for two minutes at 50% power. The lysate was rocked for 30 minutes at 4°C. After rocking for 30 minutes the lysate was separated through centrifugation. The lysates were purified by nickel affinity chromatography using His60 Ni Resin (Takara Bio). The bead volume was calculated by multiplying the wet cell mass (g) by 0.3 mL. The column was equilibrated with 3x the bead volume of cold lysis buffer. The lysate was then added to the column, and the flow through was collected and saved. The column was washed with 50 bead volumes of cold wash buffer (10 mM $Na_2HPO_4$, 10 mM Tris, 6.0 M of GuHCl, and 0.02 M imidazole, pH 8) split into two washes consisting of 25 bead volumes each of cold wash buffer. Elutions consisting of 1 bead volume of elution buffer (10 mM $Na_2HPO_4$, 10 mM Tris, 6.0 M of GuHCl, and 20 mM $CH_3COOH$, pH 3) were taken until the absorbances of each elution began to go down (usually between 6-10 elutions). Elutions were dialyzed into low salt buffer (10 mM Tris, pH 8) at 4°C. The dialysis buffer was replaced three times. The samples were further purified with anion exchange chromatography (Cytiva Q Sepharose™ Fast Flow Chromatography Media). All protein elutions were combined and centrifuged. The column was washed with low salt buffer twice to equilibrate. Twenty elutions were taken with NaCl concentrations ranging from 0 mM to 950 mM. The AbpSH3 domain eluted between 0 and 400 mM NaCl and was dialyzed into 10 mM Tris, pH 8.1 buffer for NMR.

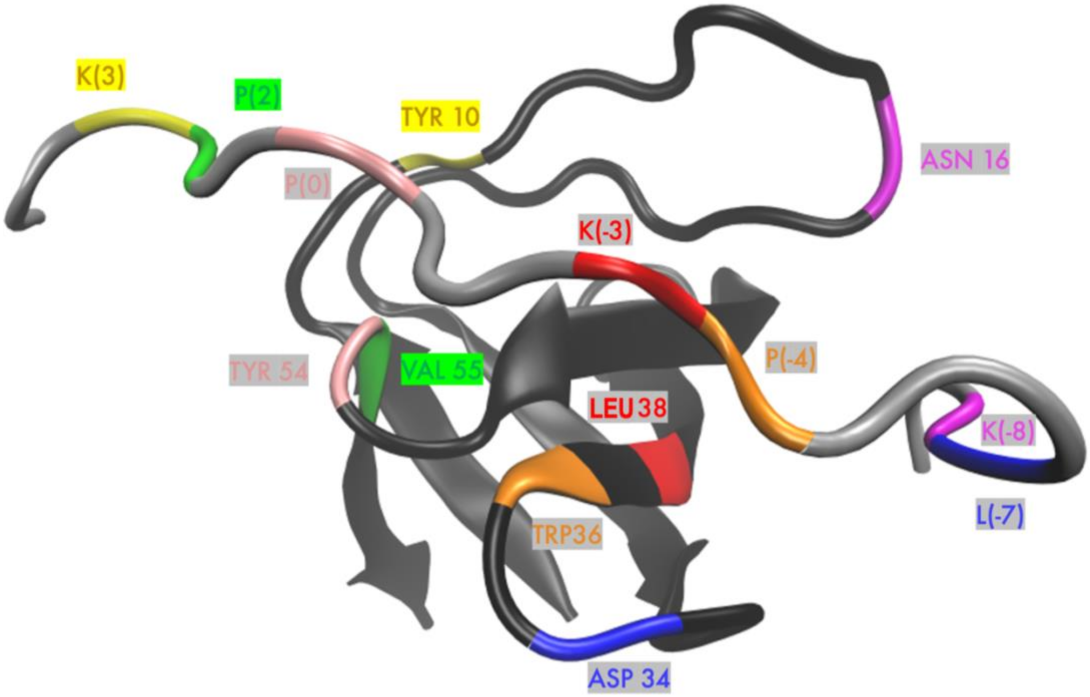


**Figure S1.** Pairwise distances used in the determination of whether the complex is in the unbound state, encounter complex, or bound state in the MD simulations. The binding surface distance was calculated as an average of the distances K(3)-Y10, P(2)-V55, P(0)-Y54, K(-3)-L38, P(-4)-W36, L(-7)-D34, and K(-8)-N16. The pairwise distance in the NMR structures of the ArkA17-AbpSH3 complex ranges from 8.73 to 8.99 Å.

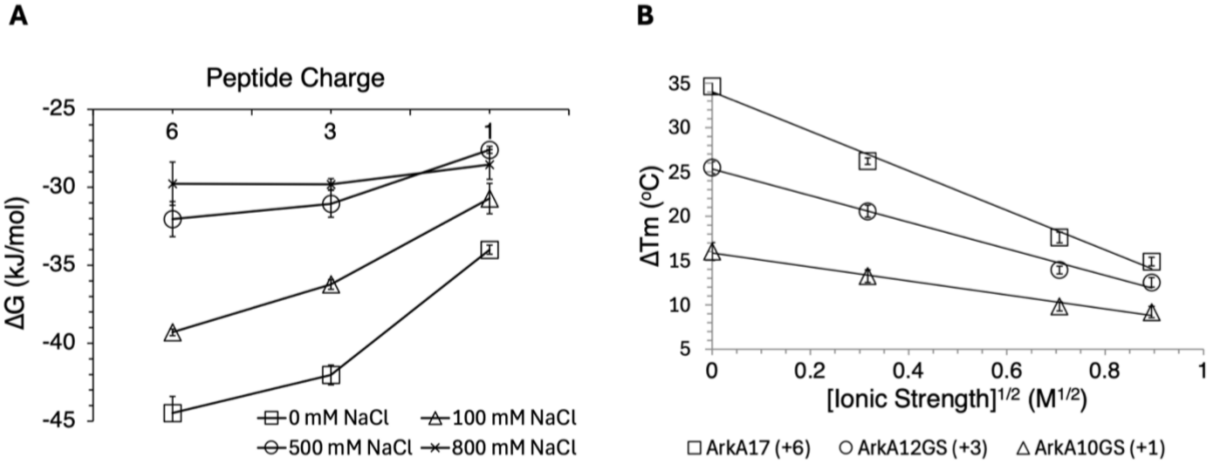


**Figure S2.** Effect of ionic strength on ArkA-SH3 complex stability. (A) Binding free energy measured by ITC plotted versus peptide charge for each salt concentration. (B) Melting temperature differences between AbpSH3-peptide hybrid and Apo AbpSH3 as measured by NanoDSF that followed tryptophan fluorescence (350 nm/330 nm emission ratio) over a temperature gradient. Data was collected at different ionic strengths for the ArkA17, ArkA12GS, and ArkA10GS peptides. ArkA12GS (GSGKPTPPPKPSHLKGS) contains the ArkA12 sequence with filler residues to keep the total length at 17.

**Table S1.** Thermodynamic and kinetic binding parameters for all peptides at 0 and 800 mM NaCl.

| peptide | [NaCl] (mM) | $K_D$ (µM) | $k_{off}$ ($s^{-1}$) | $k_{on}$ ($s^{-1}M^{-1}$) |
|---|---|---|---|---|
| ArkA17 | 0 | 0.0247 | 72.6 | $2.94 \times 10^9$ |
| ArkA17 | 800 | 7.91 | 267 | $3.38 \times 10^7$ |
| ArkA12 | 0 | 0.0582 | 70.3 | $1.21 \times 10^9$ |
| ArkA12 | 800 | 7.51 | 290 | $3.86 \times 10^7$ |
| ArkA10GS | 0 | 1.48 | NC* | NC* |
| ArkA10GS | 800 | 12.1 | NC* | NC* |

| peptide | [NaCl] (mM) | $\Delta G_{N\to B}$ (kJ $mol^{-1}$) | $\Delta\Delta G^{salt}_{N\to B}$ (kJ $mol^{-1}$) | $\Delta\Delta G^{salt}_{N\to\dagger}$ (kJ $mol^{-1}$) | $\Delta\Delta G^{salt}_{B\to\dagger}$ (kJ $mol^{-1}$) | $\Phi^{salt}_{a}$ | $\Delta\Delta G^{trunc}_{N\to B}$ (kJ $mol^{-1}$) | $\Delta\Delta G^{trunc}_{N\to\dagger}$ (kJ $mol^{-1}$) | $\Delta\Delta G^{trunc}_{B\to\dagger}$ (kJ $mol^{-1}$) |
|---|---|---|---|---|---|---|---|---|---|
| ArkA17 | 0 | -44.1 | 14.5 | 11.3 | -3.3 | 0.77 | | | |
| ArkA17 | 800 | -29.6 | | | | | | | |
| ArkA12 | 0 | -42.0 | 12.2 | 8.7 | -3.6 | 0.71 | 2.1 | 2.2 | 0.1 |
| ArkA12 | 800 | -29.7 | | | | | -0.1 | -0.3 | -0.2 |
| ArkA10GS | 0 | -34.0 | NC* | NC* | NC* | NC* | 9.4 | NC* | NC* |
| ArkA10GS | 800 | -28.5 | | | | | 0.6 | NC* | NC* |

The superscript *salt* indicates a change in $\Delta G$ between 0 and 800 mM salt concentration, while the superscript *trunc* indicates a change in $\Delta G$ between ArkA17 and either ArkA10GS or ArkA12 binding.

*For ArkA10GS, binding kinetics data was not collected.

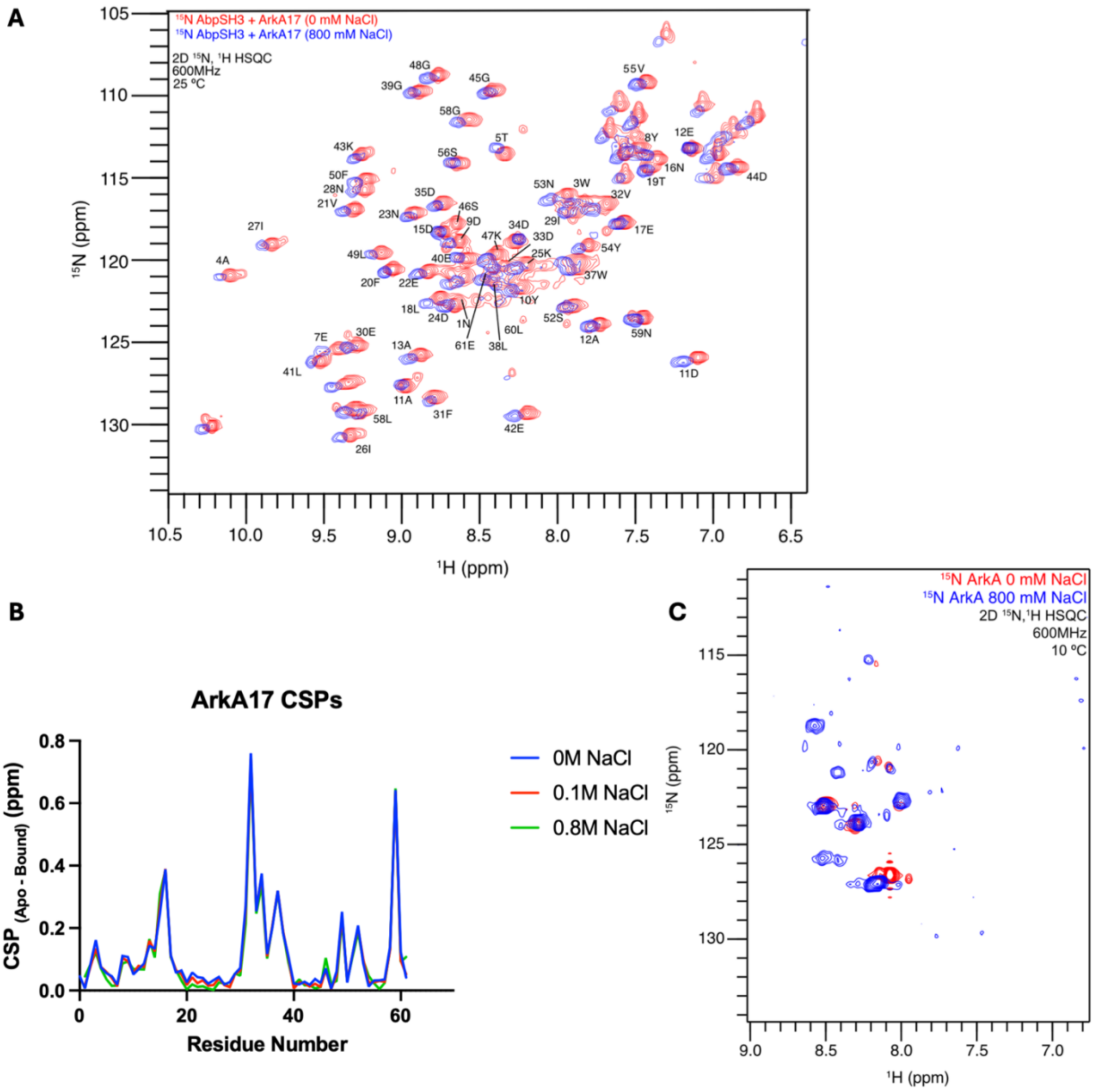


**Figure S3.** NMR HSQC data. (A) HSQC overlay of $^{15}N$-labeled AbpSH3 domain bound to ArkA17 at 0 mM (red) and 800 mM (blue) NaCl. (B) Chemical shift perturbation (CSP) plotted versus residue for the $^{15}N$-labeled AbpSH3 domain bound to ArkA17 for each NaCl concentration: 0 mM (blue), 100 mM (red), 800 mM (green). (C) HSQC overlay for free $^{15}N$-labeled ArkA17 at 0 mM (red) and 800 mM (blue) NaCl (unassigned).

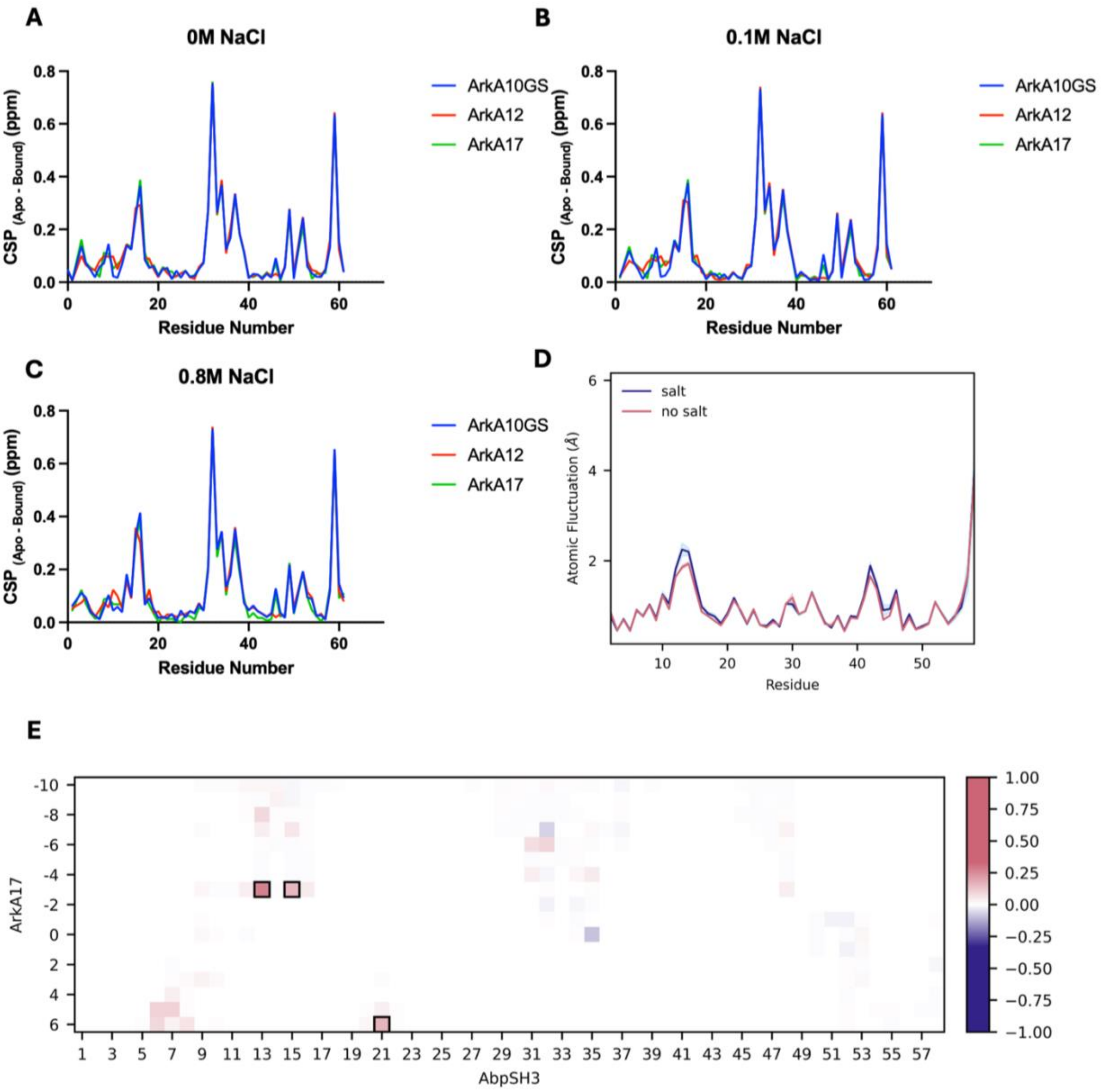


**Figure S4.** Effect of salt on the ArkA-AbpSH3 bound complex. Chemical shift perturbation (CSP) plotted versus residue for the $^{15}$N-labeled AbpSH3 domain bound to ArkA17 (green), ArkA12 (red), and ArkA10GS (blue) at (A) 0 mM NaCl, (B) 100 mM NaCl, and (C) 800 mM NaCl. (D) Root-mean-squared back bone fluctuations of AbpSH3 bound to ArkA17 from simulations at 0 mM (pink) and 900 mM (blue) salt concentrations. (E) Contact difference map from simulations of the ArkA17-AbpSH3 bound complex with and without salt. Pink contacts indicate those that are occupied more frequently in the simulations without salt and blue contacts indicate those that are occupied more frequently in simulations with 900-mM NaCl present. Boxed contacts indicate differences that have a p-value below 0.05.

**Table S2.** Overall contact map difference for ArkA17-AbpSH3 and ArkA12-AbpSH3 bound complexes and ArkA12-AbpSH3 encounter complex.

| **Construct** | **Average # of contacts no salt** | **Average number of contacts high salt** | **Difference in average number of contacts with salt** | **Sum of squares of contact population differences with salt** |
|---|---|---|---|---|
| **ArkA12 bound** | $26.11 \pm 0.31$ | $25.60 \pm 0.36$ | 0.51<br>$p = 0.1478$ | 0.097<br>$p = 0.6931$ |
| **ArkA12 encounter** | $15.89 \pm 0.44$ | $14.06 \pm 0.38$ | 1.83<br>$p = 0.002$ | 1.997<br>$p = 0.0609$ |
| **ArkA17 bound** | $30.26 \pm 0.23$ | $28.79 \pm 0.26$ | 1.47<br>$p = 0.0002$ | 1.146<br>$p = 0.0200$ |

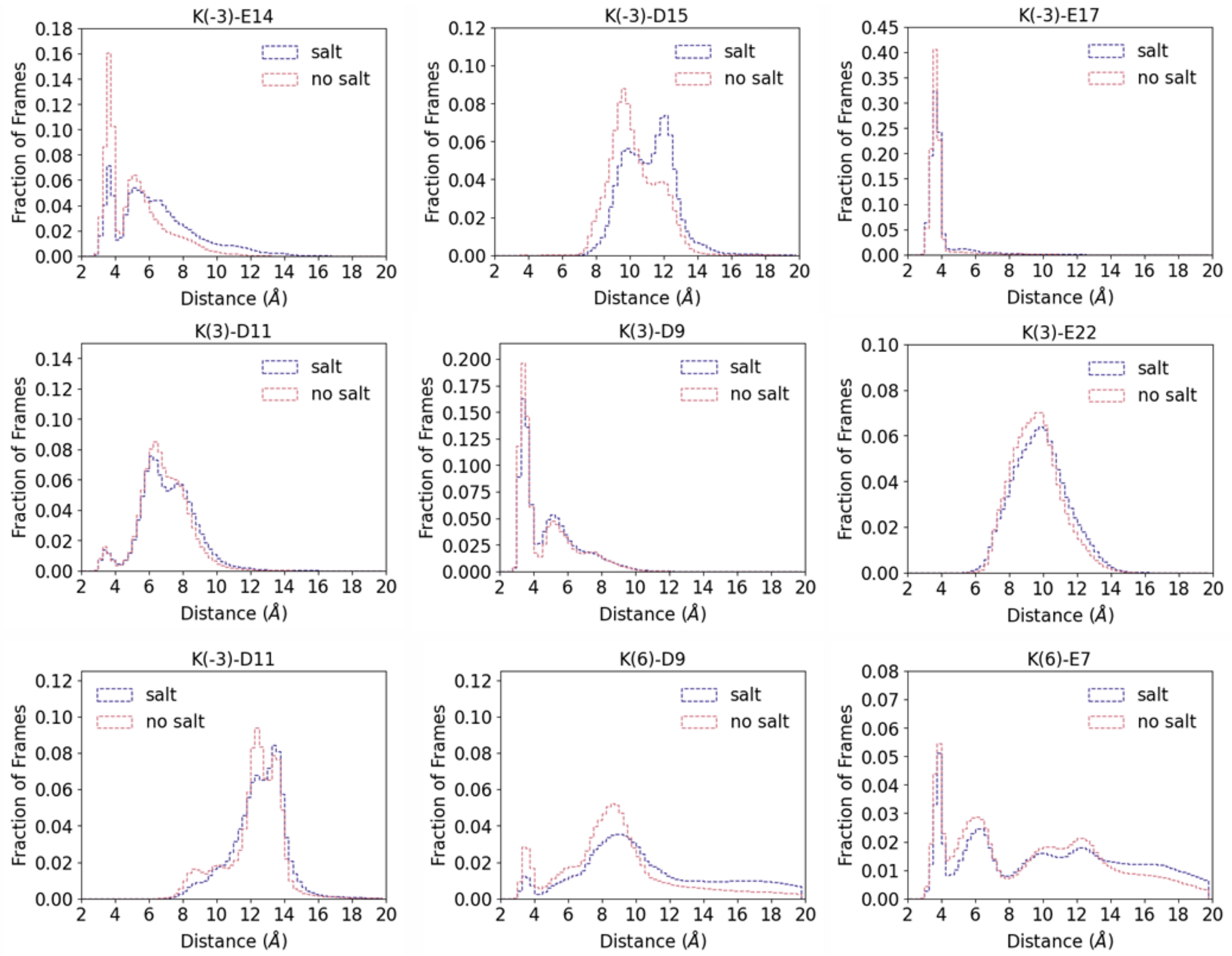

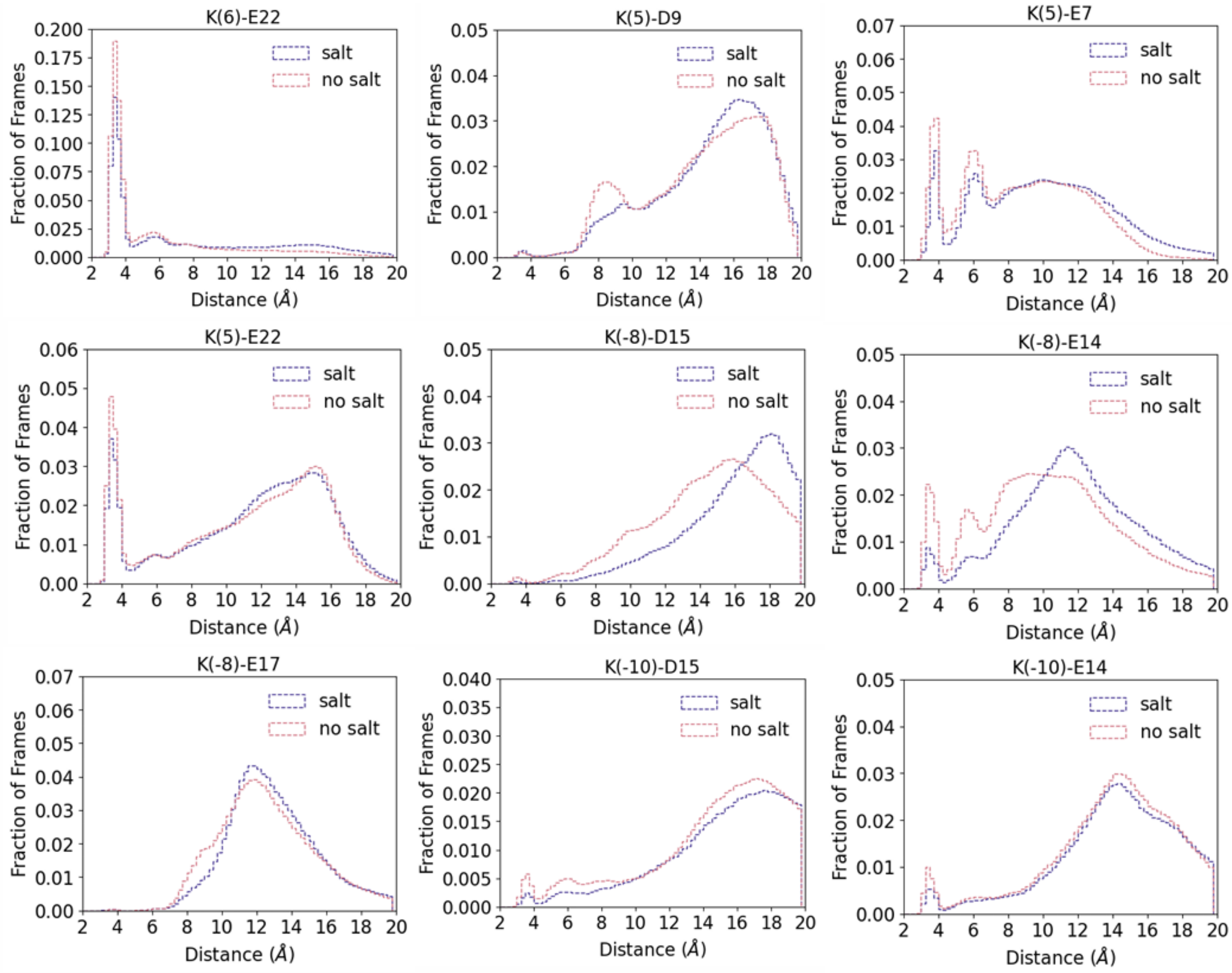


**Figure S5.** Individual ArkA17-AbpSH3 electrostatic interaction histograms from the bound complex with 0 mM and 900 mM NaCl for 18 different intermolecular pairs of charged residues. Population of the interactions are plotted versus distance for simulations at 0 mM (pink) and 800 mM (blue) salt. Distances less than 4 Å are considered short-range interactions and distances between 4 Å and 10 Å are considered long-range interactions.

**Table S3.** Statistical comparison of rate constants calculated from MD simulations using Bayesian inference.

| | $\boldsymbol{k_1}$ | $\boldsymbol{k_{on}}$ | $\boldsymbol{k_{-1}}$ |
|---|---|---|---|
| **0 mM NaCl rate constant from MD** | $5.08 \pm 0.72 \times 10^9$ $(\mathrm{s^{-1}M^{-1}})$ | $5.3 \pm 1.8 \times 10^7$ $(\mathrm{s^{-1}M^{-1}})$ | $2.54 \pm 0.42 \times 10^6$ $(\mathrm{s^{-1}})$ |
| **800 mM NaCl rate constant from MD** | $1.41 \pm 0.20 \times 10^9$ $(\mathrm{s^{-1}M^{-1}})$ | $5.3 \pm 1.8 \times 10^7$ $(\mathrm{s^{-1}M^{-1}})$ | $5.10 \pm 0.78 \times 10^6$ $(\mathrm{s^{-1}})$ |
| **Odds ratio*** | 20,160,879 | 0.381 | 17.3 |

*The odds ratio can be interpreted as follows: 0.3-3 = uncertain if there is an effect of salt; 3-10 = substantial evidence that there is an effect; greater than 10 = strong evidence that there is an effect[30].

**Table S4.** Comparison of pseudo-dissociation constant, $K[EC]DKD[EC]$, for the encounter complex using two different methods.

| | $\boldsymbol{K_D^{[EC]} = \frac{k_{-1}}{k_1}}$ | $\boldsymbol{K_D^{[EC]} = \frac{\Theta^2_{unbound}}{\Theta_{EC}}}$ |
|---|---|---|
| **0 mM NaCl** | $(0.50 \pm 0.11) \times 10^{-3}$ (M) | $(1.43 \pm 3.15) \times 10^{-5}$ (M) |
| **800 mM NaCl** | $(3.62 \pm 0.75) \times 10^{-3}$ (M) | $(1.71 \pm 3.36) \times 10^{-4}$ (M) |

**Table S5.** Differences and *p*-values comparing the average number of electrostatic interactions at different salt concentrations for the ArkA17-AbpSH3 bound complex, the ArkA12-AbpSH3 bound complex, and the ArkA12-AbpSH3 encounter complex.

| **Construct** | **No salt long** | **No salt short** | **High salt long** | **High salt short** | **Long difference** | **Short difference** | **Overall difference (no salt – high salt)** |
|---|---|---|---|---|---|---|---|
| **ArkA12 bound** | $4.17 \pm 0.04$ | $2.14 \pm 0.07$ | $4.09 \pm 0.04$ | $1.66 \pm 0.07$ | 0.08 $p = 0.0819$ | 0.48 $p = 0.0002$ | 0.56 $p = 0.0005$ |
| **ArkA12 encounter** | $4.96 \pm 0.56$ | $1.24 \pm 0.20$ | $2.41 \pm 0.38$ | $0.58 \pm 0.22$ | 2.55 $p = 0.0001$ | 0.65 $p = 0.0085$ | 3.20 $p = 0.0003$ |
| **ArkA17 bound** | $6.85 \pm 0.07$ | $2.95 \pm 0.05$ | $5.96 \pm 0.11$ | $2.26 \pm 0.05$ | 0.89 $p = 0.0001$ | 0.69 $p = 0.0001$ | 1.58 $p = 0.0001$ |

**Table S6.** Differences and *p*-values comparing number of hydrogen bonds at different salt concentrations for the ArkA17-AbpSH3 bound complex, the ArkA12-AbpSH3 bound complex, and the ArkA12-AbpSH3 encounter complex.

| **Construct** | **Hydrogen bonds at no salt** | **Hydrogen bonds at high salt** | **Overall difference (no salt – high salt)** |
|---|---|---|---|
| **ArkA12 bound** | $5.03 \pm 0.12$ | $4.44 \pm 0.13$ | 0.59 $p = 0.0023$ |
| **ArkA12 encounter** | $2.91 \pm 0.12$ | $1.96 \pm 0.10$ | 0.86 $p = 0.0001$ |
| **ArkA17 bound** | $6.27 \pm 0.08$ | $5.28 \pm 0.08$ | 0.99 $p = 0.0001$ |

**Table S7.** Differences and *p*-values comparing solvent accessible surface area (SASA) at different salt concentrations for the ArkA12-AbpSH3 bound complex and the ArkA12-AbpSH3 encounter complex.

| Construct | SASA at no salt | SASA at high salt | Overall difference (no salt – high salt) |
|---|---|---|---|
| **ArkA12 bound** | $39.925 \pm 0.001$ | $39.930 \pm 0.001$ | 0.25<br>$p = 0.6155$ |
| **ArkA12 encounter** | $44.744 \pm 0.001$ | $45.380 \pm 0.001$ | 0.64<br>$p = 0.7784$ |

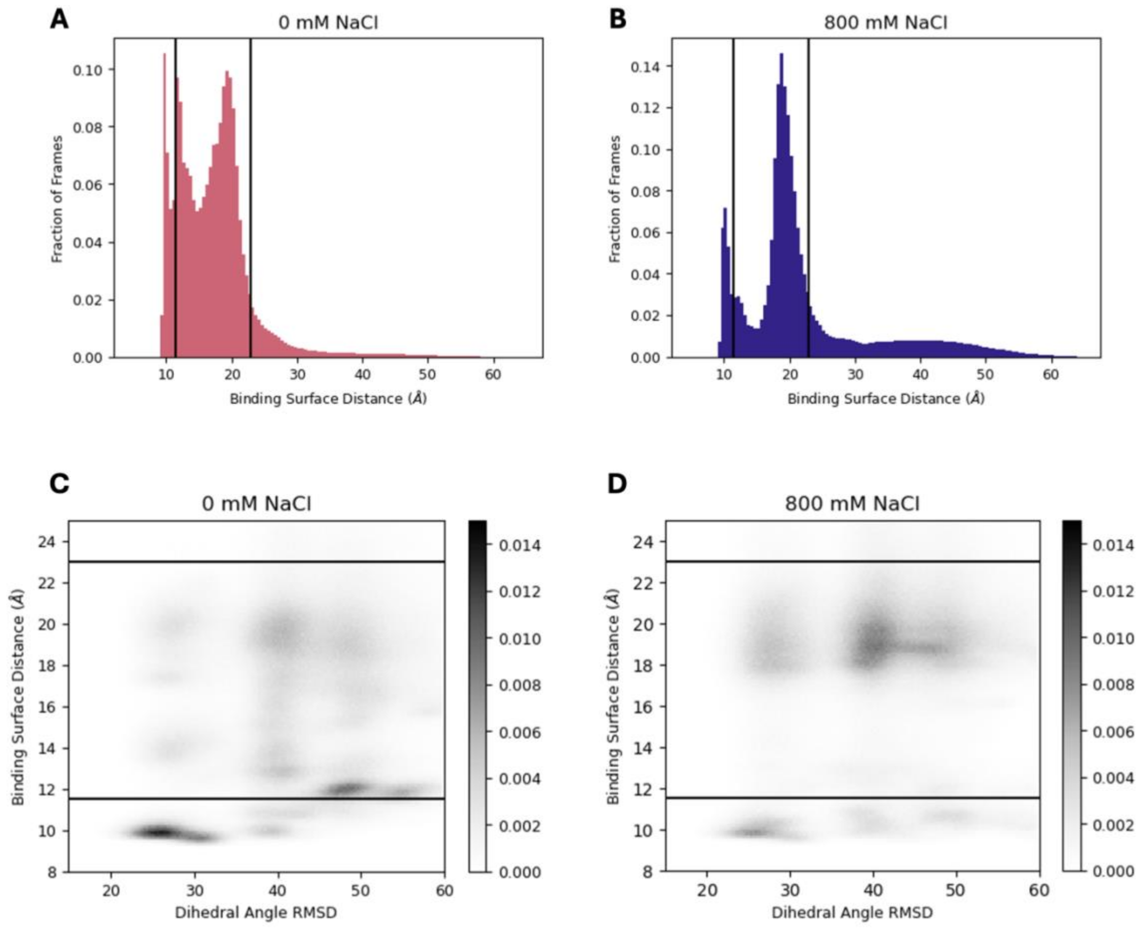


**Figure S6.** Population histogram of binding surface distance from ArkA12 binding in (A) 0 mM salt and (B) 800 mM salt. States left of the vertical line at 11.5 Å are classified as bound. States between the vertical lines at 11.5 Å and 23 Å are classified as the encounter comples. States to the right of the vertical line at 23 Å are classified as unbound. Binding surface distance plotted versus ArkA backbone dihedral angle RMSD for the simulations of ArkA12 binding to AbpSH3 at (C) 0 mM NaCl and (D) 800 mM NaCl. Darker shading indicates a larger fraction of the total ensemble, as indicated by the color bar.

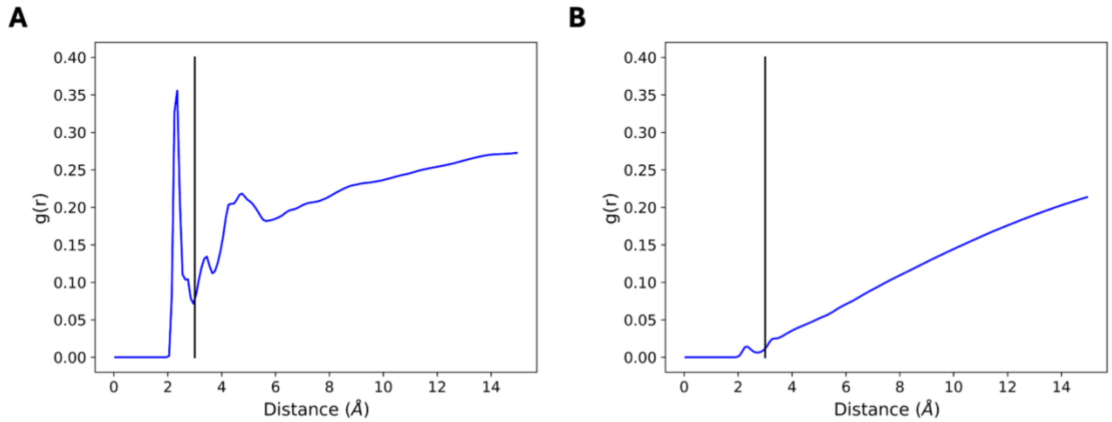


**Figure S7.** Radial distribution functions from simulations of the ArkA17-AbpSH3 bound complex in 900 mM NaCl showing the typical ion contact distances for (A) $Na^+$ ions and (B) $Cl^-$ ions. The vertical line at 3 Å shows our cutoff for defining an ion contact used in Table 5.

**Table S8.** Number of unique ions interacting with each residue during a single simulation.

| **Residue** | **Apo** | **ArkA12 EC** | **ArkA12 Bound** |
|---|---|---|---|
| Glu7 | $50.8 \pm 0.13$ | $150.24 \pm 5.13$ | $52.0 \pm 0.00$ |
| Asp9 | $51.0 \pm 0.00$ | $127.98 \pm 5.65$ | $52.0 \pm 0.00$ |
| Asp11 | $51.0 \pm 0.00$ | $142.4 \pm 5.95$ | $52.0 \pm 0.00$ |
| Glu14 | $50.8 \pm 0.13$ | $154.04 \pm 5.61$ | $52.0 \pm 0.00$ |
| Asp15 | $50.8 \pm 0.13$ | $165.94 \pm 5.21$ | $52.0 \pm 0.00$ |
| Glu17 | $50.8 \pm 0.13$ | $66.18 \pm 5.84$ | $35.0 \pm 1.06$ |
| Glu22 | $50.8 \pm 0.13$ | $153.3 \pm 5.22$ | $52.0 \pm 0.00$ |
| Asp24 | $49.5 \pm 0.40$ | $36.12 \pm 2.35$ | $48.8 \pm 0.61$ |
| Lys25 | $39.0 \pm 0.00$ | $161.32 \pm 5.61$ | $43.0 \pm 0.00$ |
| Glu30 | $50.8 \pm 0.13$ | $143.84 \pm 5.57$ | $52.0 \pm 0.00$ |
| Asp33 | $50.8 \pm 0.13$ | $128.38 \pm 5.71$ | $52.0 \pm 0.00$ |
| Asp34 | $50.8 \pm 0.13$ | $156.94 \pm 5.14$ | $52.0 \pm 0.00$ |
| Asp35 | $50.8 \pm 0.13$ | $137.22 \pm 5.25$ | $52.0 \pm 0.00$ |
| Glu40 | $50.8 \pm 0.13$ | $93.76 \pm 4.54$ | $52.0 \pm 0.00$ |
| Glu42 | $50.8 \pm 0.13$ | $136.18 \pm 5.47$ | $51.9 \pm 0.10$ |
| Lys43 | $39.0 \pm 0.00$ | $159.52 \pm 6.05$ | $43.0 \pm 0.00$ |
| Asp44 | $50.8 \pm 0.13$ | $120.84 \pm 5.86$ | $52.0 \pm 0.00$ |
| Lys47 | $39.0 \pm 0.00$ | $169.18 \pm 4.57$ | $43.0 \pm 0.00$ |
